# Widespread cryptic RAG-mediated recombination in developing T lymphocytes

**DOI:** 10.64898/2026.09.01.748595

**Authors:** D. Sobral, J. Silva, D. Santos, J. Perreira Leal, S. Spicuglia, J. Demengeot, M. Bonnet-Di Placido

## Abstract

V(D)J recombination generates antigen receptor diversity through the targeted activity of the RAG1/2 recombinase, but the extent to which RAG also engages cryptic genomic targets during normal lymphocyte development remains poorly defined. Here, we developed a targeted PCR-sequencing approach to detect and quantify rare RAG-mediated rearrangements in mouse thymocytes. We first examined the TCRβ locus and found that four of its twelve pseudogenes undergo detectable rearrangement in vivo, despite being considered non-functional components of the repertoire. Extending this analysis across the locus revealed 33 previously uncharacterized cryptic recombination sites involved in non-functional rearrangements with DJβ segments. These events occurred across both the Vβ region and the largely inaccessible Vβ30–Dβ1 intergenic region, with individual cryptic sites spanning a broad range of recombination frequencies. Cryptic sites were highly heterogeneous in sequence and chromatin context: neither RSS resemblance, predicted Z-DNA formation, local epigenetic features, nor chromosomal interactions reliably distinguished sites with detectable recombination from those at which recombination was not detected. We further identified additional cryptic RAG-mediated rearrangements at the Bcl11b locus, demonstrating that such events are not restricted to antigen receptor loci. Together, these findings reveal an unexpectedly broad landscape of low-frequency RAG-mediated DNA rearrangement in developing T lymphocytes and suggest that cryptic target selection cannot be explained solely by the local genetic and epigenetic features examined here.

## INTRODUCTION

V(D)J recombination generates the diversity of antigen receptors through the somatic assembly of variable (V), diversity (D) and joining (J) gene segments in developing lymphocytes. This process is initiated by the lymphoid-specific recombination-activating proteins RAG1 and RAG2, which recognize recombination signal sequences (RSSs) flanking antigen receptor gene segments (1). RAG-mediated cleavage generates DNA double-strand breaks at the junction between the RSS and coding sequence, which are subsequently repaired by the non-homologous end-joining machinery. Imprecise processing of the coding ends, including nucleotide deletion and the addition of palindromic and non-templated nucleotides, further contributes to antigen receptor diversity and provides a characteristic molecular signature of RAG-mediated recombination (1, 2).

Canonical RSSs consist of conserved heptamer and nonamer motifs separated by either a 12- or 23-bp spacer (12-RSS and 23-RSS, respectively), with efficient recombination generally requiring pairing of one of each type (2, 3). However, RSSs are intrinsically degenerate, and RAG can recognize a broad range of sequence variants (4). Consequently, RSS-like sequences elsewhere in the genome, referred to as cryptic RSSs (cRSSs), can also become substrates for RAG-mediated cleavage and recombination (5–9). Sequence-based algorithms, including the Recombination Information Content (RIC) score (10) and the Recombination Efficiency Coefficient (REC) score (11), have therefore been developed to predict the recombination potential of RSS-like sequences. In addition to RSS-like sequences, non-B DNA structures such as Z-DNA can be recognized by the RAG complex and participate in aberrant rearrangements (12).

Cryptic RAG activity has important implications for genome integrity. RAG-mediated rearrangements involving non-antigen receptor loci have been identified in lymphoid malignancies and can contribute to oncogenic deletions and translocations (9, 13, 14). However, cryptic recombination is not restricted to malignant cells: low-frequency RAG-dependent rearrangements have also been detected in lymphocytes from healthy individuals and mice (15–18). Individual events have been reported at frequencies reaching approximately 100 rearrangements per million cells (6–8, 19). More recently, genome-wide approaches have revealed a broader spectrum of RAG off-target activity. High-throughput analysis of the TCRδ locus identified numerous off-target RAG rearrangements generated within chromosomal loop domains in developing T cells (20), while direct genome-wide mapping of DNA double-strand breaks by END-seq demonstrated RAG-dependent cleavage at cryptic sites, including in primary thymocytes (21). Nevertheless, most studies have focused on cleavage events, selected rearrangements or genomic lesions with functional or pathological consequences. The frequency, sequence diversity and local determinants governing low-frequency cryptic rearrangements during normal lymphocyte development therefore remain incompletely understood.

RAG targeting is determined not only by RSS sequence and local chromatin accessibility, but also by the three-dimensional organization of antigen receptor loci. Regions undergoing V(D)J recombination are enriched for features associated with accessible chromatin, including H3K4me3, histone acetylation, RNA polymerase II occupancy and germline transcription (22, 23). RAG2 directly recognizes H3K4me3 through its plant homeodomain (24, 25), while RAG1 occupancy is associated with acetylated chromatin (24). Consistent with this, epigenetic features associated with chromatin accessibility correlate with gene-segment usage at the IgH and TCRβ loci (26, 27). Subsequent studies have further established that RAG bound at recombination centres can scan chromatin for potential substrates and that this process is strongly influenced by CTCF-binding elements and cohesin-mediated chromatin loop extrusion (28–30). RSS orientation is an important determinant of this scanning process, which can involve both canonical and cryptic RSSs (20, 30). Thus, RAG target selection reflects the interplay between RSS sequence, local chromatin environment and higher-order locus architecture. How these features determine which cryptic sites ultimately undergo productive DNA rearrangement in vivo, however, remains poorly understood.

The mouse TCRβ locus provides a particularly informative system in which to address this question. V(D)J recombination at this locus occurs during the double-negative stages of thymocyte development and has been extensively characterized (31, 32). The ∼700-kb locus contains 22 functional Vβ gene segments and 12 Vβ pseudogenes, followed by two Dβ-Jβ-Cβ clusters and the isolated Vβ30 gene segment. Importantly, these regions occur within markedly different chromatin environments: the Vβ and Dβ-Jβ-Cβ regions are accessible during recombination, whereas the large Vβ30-Dβ1 intergenic region is predominantly associated with closed chromatin (27, 33–38). Moreover, Vβ-associated RSSs vary considerably in sequence and recombination efficiency, contributing to the non-uniform usage of Vβ gene segments in the TCRβ repertoire (39–42). Previous repertoire studies have also provided evidence that non-functional Vβ pseudogenes can undergo rearrangement (27, 43), suggesting that the locus may contain additional, poorly characterized RAG substrates.

Here, we developed a targeted genomic PCR-sequencing approach to detect rare RAG-mediated rearrangements in mouse thymocytes and used the TCRβ locus to investigate the prevalence and determinants of cryptic recombination. We first examined Vβ pseudogene usage and subsequently extended the analysis to cryptic sites across regions with contrasting chromatin accessibility. We identified a broad set of previously uncharacterized cryptic rearrangements, including events within the predominantly inaccessible Vβ30-Dβ1 region, and further detected additional cryptic RAG-mediated rearrangements at the Bcl11b locus. By integrating sequence characteristics, predicted Z-DNA formation, epigenetic features and chromosomal interactions, we then asked whether these properties could distinguish sites with detectable cryptic recombination from those at which recombination was not detected. Together, our findings reveal a broad landscape of low-frequency RAG-mediated DNA rearrangement during normal T-cell development and show that cryptic target selection cannot be explained solely by local RSS sequence and epigenetic features.

## RESULTS

### Vβ pseudogene rearrangements reveal cryptic RAG-mediated recombination at the TCRβ locus

Among the 12 Vβ pseudogenes at the TCRβ locus, all except Vβ10 contain at least one stop codon, four lack a leader sequence (Vβ6, Vβ8, Vβ9 and Vβ10), at least four have inactive promoters (Vβ7, Vβ11, Vβ22 and Vβ28; (27)), and seven are flanked by RSSs with REC scores below the recombination thresholds (Vβ8, Vβ11, Vβ12.3, Vβ18, Vβ25, Vβ27 and Vβ28). To our knowledge, only two previous studies have assessed pseudogene usage at the mouse TCRβ locus (27, 43). Overall, the repertoires reported in these studies correlate relatively well (R²=0.74, Figure S1). However, only one rearranging pseudogene was detected in one study (27), whereas the other reported evidence of *in vivo* recombination for all pseudogenes (43).

To assess TCRβ pseudogene usage more directly, we developed three complementary targeted approaches based on (semi-)nested PCR followed by sequencing or Southern blotting (Figure 1). First, we performed semi-nested PCR on five-fold serial dilutions of WT genomic DNA (gDNA) from total thymocytes and early DN3 cells (DN3E; before β-selection). Primers were designed to amplify rearrangements of the Vβ22, Vβ10 and Vβ6 pseudogenes with the DJβ1 or DJβ2 clusters (43) (see Methods and Table S1A). Rearrangements of Vβ22 and Vβ10 with the Jβ2 (Figure 2A) and Jβ1 (Figure S2A) clusters were detected, with Vβ22 remaining detectable one dilution further than Vβ10. Similar patterns were observed in total thymocytes and DN3E cells, consistent with the limited effect of β-selection on relative Vβ usage previously reported (44, 45) (correlation coefficients of 0.97 and 0.90 before and after β-selection, respectively; Figure S1). For Vβ6, a single band was detected following amplification of DN3E gDNA with the Jβ2.7 reverse primer. Unexpectedly, sequencing revealed that this amplicon did not correspond to Vβ6 rearrangement, but instead to recombination of Jβ2.2 with a cryptic site located approximately 300 bp downstream of Vβ6, hereafter designated cV6. The coding joint displayed characteristic features of RAG/NHEJ-mediated recombination, including nucleotide deletion and addition, CG enrichment and palindromic nucleotides (Table 1A).

**Figure 1.**
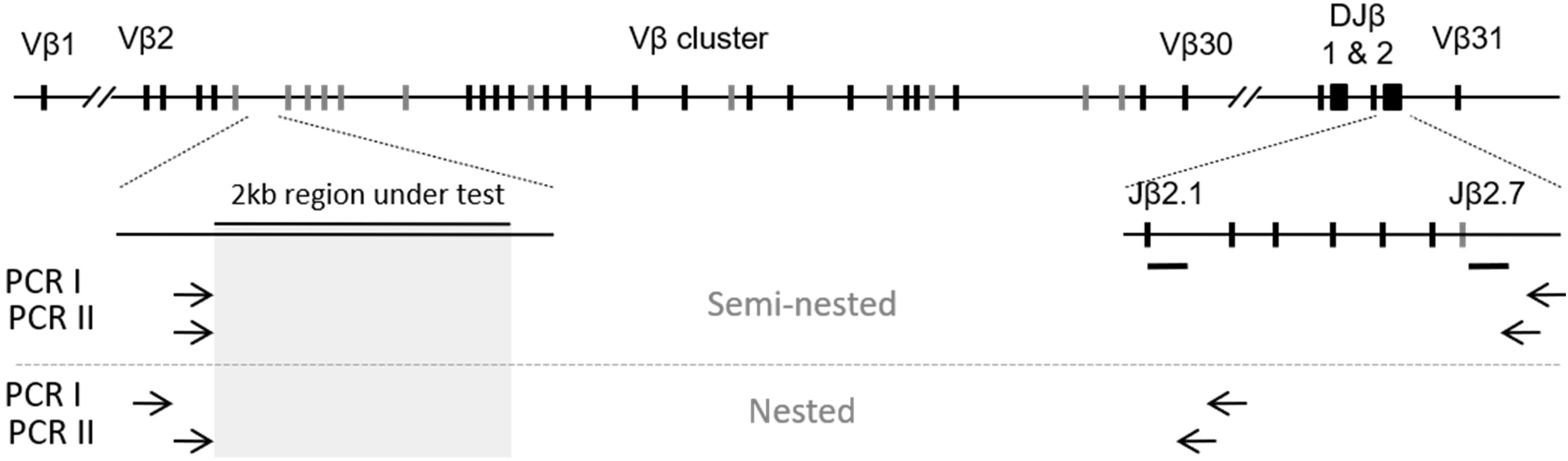
Amplification strategy for cRSS detection at the TCRβ locus. The top schematic represents the structure of the TCRβ locus, with functional gene segments and pseudogenes indicated by black and grey vertical lines, respectively. Long intergenic regions are not shown to scale, including those between Vβ1 and Vβ2 and between Vβ30 and Dβ1. Enlarged schematics of a representative region tested for cRSSs within the Vβ region and of the Jβ2 cluster illustrate the positions of primers used for semi-nested PCR (above the dotted line) and nested PCR (below the dotted line). The positions of the radioactive Jβ-specific probes are indicated by horizontal lines.

**Figure 2.**
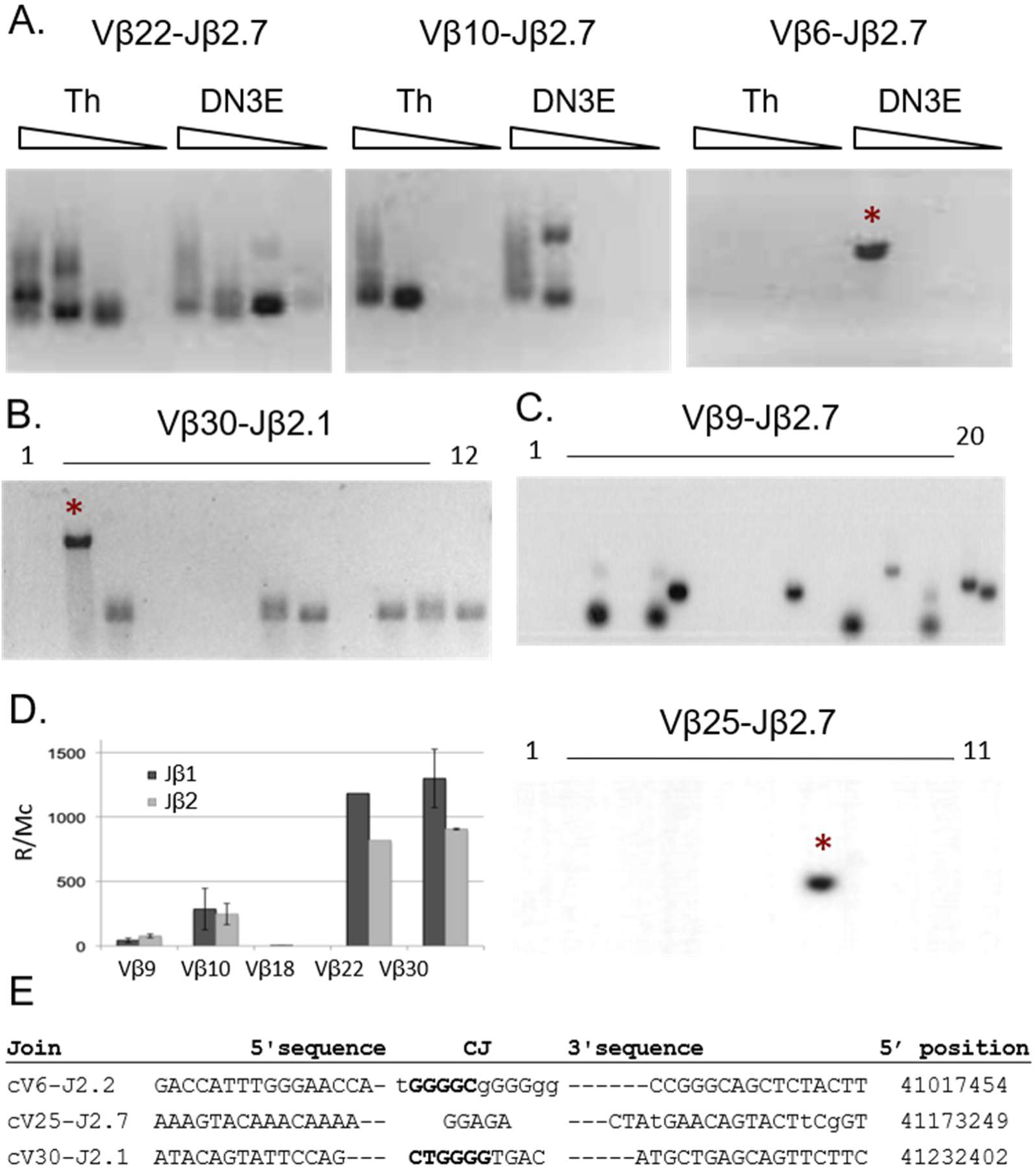
Detection and quantification of low-frequency Vβ pseudogene and cryptic rearrangements. (A) Agarose gel images of PCR amplification of five-fold serial dilutions of WT thymocyte gDNA, from 100 ng to 0.8 ng per reaction, showing Vβ22, Vβ10 and Vβ6 rearrangements with Jβ2.7. (B) Agarose gel image of 12 PCR reactions amplifying Vβ30–Jβ2.1 rearrangements, using 20 ng of gDNA per reaction. (C) Autoradiograph of Southern blot analysis of PCR reactions amplifying Vβ9–Jβ2.7 and Vβ25–Jβ2.7 rearrangements, using 60 ng of WT thymocyte gDNA per reaction. (D) Estimated rearrangement frequencies of the detected Vβ pseudogene segments, calculated from fluctuation-PCR analyses. (E) Coding-joint sequences of cryptic rearrangements identified by PCR. Asterisks (*) indicate cryptic rearrangement products.

**Table 1.**
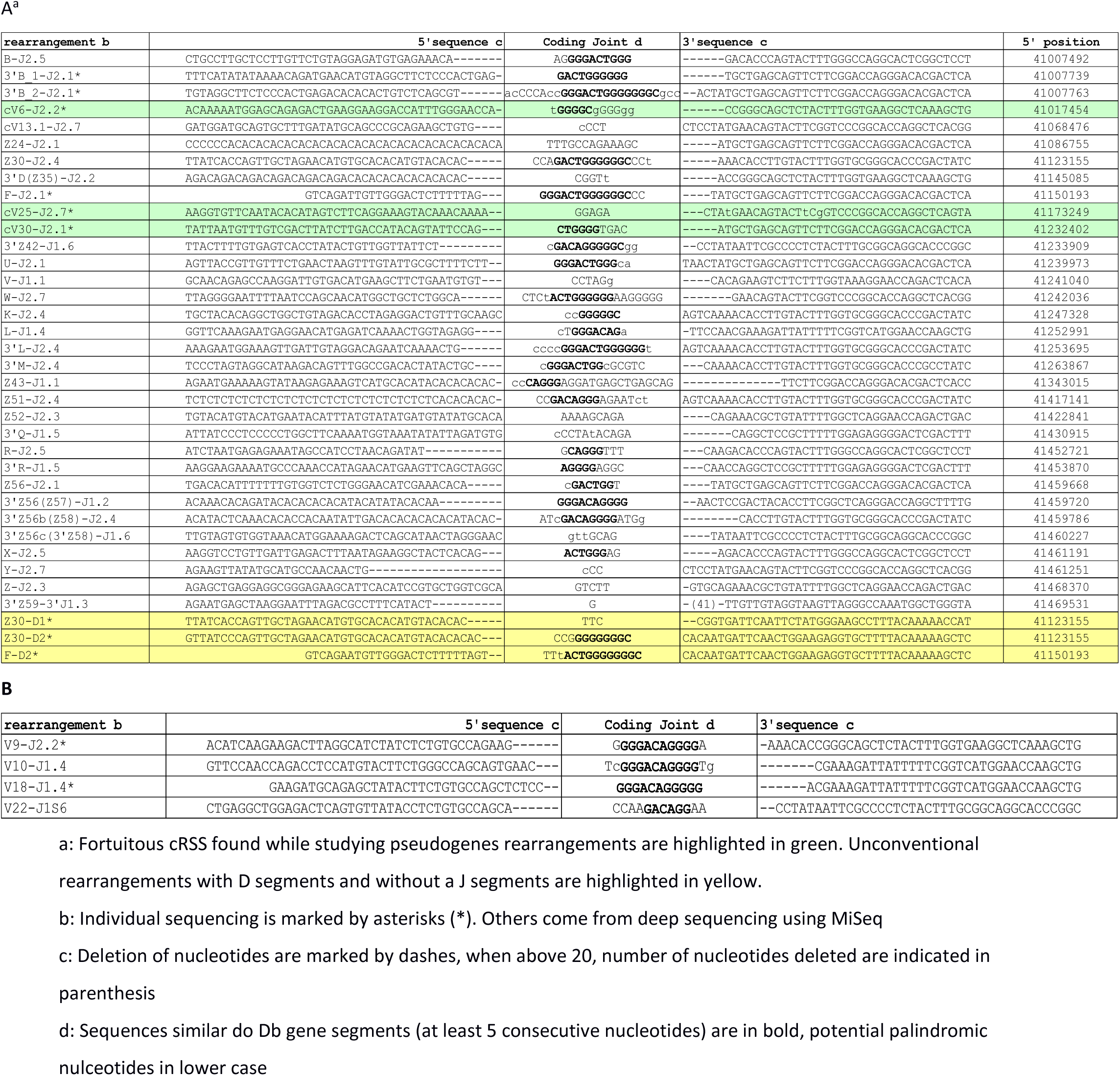
Examples of cryptic sites (A) and pseudogenes (B) rearrangements coding joints.

Second, we used semi-nested fluctuation PCR, repeating individual PCR reactions as previously described (16), to obtain approximate estimates of pseudogene rearrangement frequencies. All 12 pseudogenes were tested for recombination with both complete DJβ clusters using total-thymocyte gDNA from WT mice, with RAG2-KO thymocyte gDNA as a negative control (see Methods, (27) and Table S1A). For each pseudogene, 2,640 ng of gDNA from each genotype, corresponding to approximately 440,000 cells, was analysed. As a reference, Vβ30 rearrangements with the first gene segment of each cluster (Jβ1.1 and Jβ2.1) were also quantified. We detected 16 Vβ30 rearrangements in 480 ng of gDNA. Assuming equivalent usage of the individual Jβ gene segments (27, 46), this corresponds to an estimated Vβ30 rearrangement frequency of approximately 2,200 rearrangements per million cells (R/Mc) (Figures 2B and S2B). An additional band of unexpected size was detected in the Vβ30-Jβ2.1 assay (Figure 2B). Sequencing identified this product as a rearrangement between DJβ2.1 and a cryptic site approximately 400 bp downstream of Vβ30, hereafter cV30, again displaying features characteristic of RAG/NHEJ-mediated recombination (Table 1A). No bands were detected for Vβ27 or Vβ28 (data not shown), whereas several other pseudogene assays generated products in both WT and RAG2-KO samples.

Third, to distinguish specific rearrangement products from non-specific amplification, PCR products were analysed by Southern blotting using Jβ1.6- or Jβ2.7-specific radioactive probes. No hybridising bands were detected for Vβ6, Vβ7, Vβ8 or Vβ11 with either the Jβ1 or Jβ2 cluster (data not shown), indicating that, if these pseudogenes rearrange, their frequency was below the detection limit of approximately 2.3 R/Mc. In contrast, at least one hybridising band was detected for all other pseudogenes tested using WT gDNA (Figures 2C and S2C,D), whereas no hybridising bands were observed in any RAG2-KO sample. Positive amplicons were subsequently pooled and deep-sequenced to confirm the identity of the rearrangements.

Sequencing confirmed Vβ10 and Vβ22 rearrangements (Tables 1B and S2), with estimated frequencies of 304 R/Mc for Vβ10 (7 rearrangements in 138 ng; Figure S2C) and 1,182 R/Mc for Vβ22 (13 rearrangements in 66 ng; Figure S2C) (Figure 2D). Rearrangements of Vβ10 and Vβ22 with the pseudogene Jβ1.6 were also detected, demonstrating usage of pseudogene Jβ segments (Table S2). Vβ9 rearrangements occurred at an estimated frequency of approximately 60 R/Mc (12 rearrangements in 1,200 ng; Figures 2C,D). For Vβ18, a single rearrangement was detected among 2,640 ng of gDNA and confirmed by sequencing as Vβ18-Jβ1.4 (Figure S2D and Table 1B), corresponding to approximately 2.3 R/Mc (Figure 2D). The particularly low frequency of Vβ18 rearrangement may reflect the low predicted quality of its associated RSS, which does not pass the REC sensitivity threshold. Numerous bands were detected for Vβ12.3 (Figure S2D), but sequencing reads aligned preferentially to other Vβ12 gene segments, indicating insufficient primer specificity; Vβ12.3 was therefore excluded from further analysis. Finally, the Vβ25 Southern blot contained a single positive amplicon that sequencing identified as rearrangement of Jβ2.7 with a cryptic site approximately 1 kb downstream of Vβ25, hereafter cV25 (Figure 2C and Table 1A).

Overall, rearrangement was directly confirmed for four Vβ pseudogenes-Vβ18, Vβ9, Vβ10 and Vβ22-at frequencies ranging from approximately 2.3 to 1,182 R/Mc. We did not confirm rearrangement of Vβ8, despite this being the most frequently used pseudogene reported by Li *et al.* (43). Conversely, Vβ22 rearranged at the same order of magnitude as the functional Vβ30 gene segment (1,182 versus 2,200 R/Mc; Figure 2D), consistent with previous observations (27, 43). Thus, semi-nested fluctuation PCR combined with Southern blotting and deep sequencing enabled detection of rare non-functional TCRβ rearrangements down to an approximate frequency of 2.3 R/Mc. Importantly, this analysis also unexpectedly identified three cryptic recombination sites downstream of Vβ6, Vβ25 and Vβ30 (Figure 2E), prompting us to investigate the extent of cryptic recombination across the locus.

### Targeted analysis reveals multiple cryptic recombination sites across the Vβ region

The four recombining pseudogenes identified above provided too few sites to investigate the determinants of cryptic recombination. However, the incidental identification of three cRSSs in the vicinity of Vβ pseudogenes suggested that additional cryptic RAG substrates might be present throughout the TCRβ locus. We therefore extended our targeted PCR-sequencing strategy to search for 23-RSS-like sequences in the same orientation as the RSSs of Vβ1–Vβ30, thereby focusing on cryptic rearrangements that would mimic canonical deletional V(D)J recombination.

Because non-B DNA structures have also been implicated in RAG-mediated rearrangements (12), we additionally included predicted Z-DNA motifs. Although sequence requirements for Z-DNA-associated RAG recombination are not well defined, we prioritised Z-DNA-forming sequences containing at least one CA dinucleotide as candidate deletional recombination sites. We identified 124 predicted Z-DNA sequences across the TCRβ locus, 70 of which contained at least one CA. Fifteen candidate sites within the Vβ1–Vβ30 region were selected for experimental analysis: eight associated with predicted Z-DNA motifs (Z2, Z7, Z10, Z20, Z24, Z25, Z27 and Z30) and seven associated with favourable REC scores (A–G; Table S1A). Under the PCR conditions used, rearrangements involving cryptic sites up to approximately 2 kb downstream of the forward primer could potentially be amplified, allowing individual assays to reveal additional non-targeted cryptic sites.

Deep sequencing identified rearrangements involving seven cryptic sites and (D)Jβ gene segments (Tables 1 and S4), four of which corresponded directly to the 15 candidate sites tested. These included two REC-selected sites (B and F) and two Z-DNA-selected sites (Z24 and Z30). Three additional cryptic sites were identified incidentally: a predicted Z-DNA structure 1.1 kb downstream of candidate D, subsequently designated Z35, and two sites located 247 and 271 bp 3′ of candidate B. Coding joints from all seven sites displayed features consistent with RAG-mediated recombination (Tables 1B and S4). Notably, three products represented direct rearrangements with Dβ segments before Dβ-to-Jβ joining (Z30-Dβ1, Z30-Dβ2 and F-Dβ2; Table 1).

Thus, targeted interrogation of only 15 candidate regions identified seven cryptic recombination sites, including three that had not themselves been selected as candidates, indicating that cryptic RAG-mediated rearrangements are distributed throughout the accessible Vβ region.

### Cryptic RAG-mediated rearrangements also occur within the Vβ30–Dβ1 intergenic region

The Vβ1–Vβ30 region displays epigenetic features characteristic of accessible chromatin during TCRβ recombination (27, 33–38), potentially facilitating access of RAG not only to canonical RSSs but also to cryptic substrates. In contrast, the Vβ30– Dβ1 intergenic region is predominantly associated with a less accessible chromatin environment (34, 35, 37, 38). We therefore asked whether cryptic rearrangements could also be detected within this region.

Twenty-six candidate sites between Vβ30 and Dβ1 were analysed (Table S3). Thirteen were located upstream of predicted Z-DNA motifs (Z42–Z47, Z50–Z53, Z55, Z56 and Z59), and 13 were located upstream of potential cRSSs with favourable REC scores (H–T). Because repetitive sequences within the Vβ30–Dβ1 region complicated primer design and product assignment, the fluctuation-PCR assay was adapted from a semi-nested to a nested-PCR format.

The equivalent of one million WT thymocytes was analysed for rearrangements between each candidate region and both DJβ clusters. Half of each PCR reaction was analysed by agarose gel electrophoresis. Products were observed for all candidate sites except J, Z55, S and T (data not shown). The remaining material from PCRs containing at least one band was pooled and subjected to deep sequencing (Tables 1A and S3).

Deep sequencing confirmed 23 cryptic sites within the products generated from this region (Tables 1A and S4). Seven breakpoints were located directly downstream of the corresponding assay primers (K, L, Z43, Z51, Z52, R and Z56), while 12 were located up to 1.5 kb downstream (3′Z42, 3′H, 3′I, 3′L, 3′M, 3′Q, 3′R, five sites 3′ of Z56-Z57, Z58, 3′Z58a, 3′Z58b and 3′Z58c-and 3′Z59). For three additional rearrangements, the originating PCR could not be determined because products had been pooled before sequencing. Two of these sites were within the Vβ30–Dβ1 region and were designated U and V, whereas the third was located approximately 2 kb downstream of Vβ13.1 and was designated cVβ13.1 (Tables 1A and S4). Conversely, nine assays that produced visible bands (N, Z44, Z45, O, P, Z46, Z47, Z50 and Z53) yielded no sequence corresponding to the expected coding-joint region and were therefore considered negative.

These results demonstrate that cryptic RAG-mediated rearrangements are not confined to the accessible Vβ region but can also be detected extensively within the predominantly inaccessible Vβ30–Dβ1 intergenic region.

### Cryptic sites span a broad range of recombination frequencies

In total, 52 candidate sites, including the 12 Vβ pseudogenes, were experimentally interrogated for rearrangement with the DJβ clusters. Across these analyses, we identified 33 cryptic sites, including several sites that were detected incidentally rather than specifically targeted, while 33 sites tested by PCR showed no detectable rearrangement (Table S3). Ten cryptic sites were identified across the 27 loci tested within the Vβ region, corresponding to approximately 54 kb of effectively interrogated sequence, and 16 cryptic sites were identified downstream of the 26 loci tested within the Vβ30–Dβ1 region, corresponding to approximately 52 kb. Across the ∼106 kb effectively interrogated, 26 cryptic sites were identified, giving an observed density of approximately 0.25 cryptic 23-RSS-like sites per kb. This is approximately ten-fold lower than the theoretical density previously predicted from sequence alone (5). A simple extrapolation of this observed density across the ∼700-kb TCRβ locus would correspond to approximately 172 such sites; however, because candidate regions were selected rather than randomly sampled, this value should be considered illustrative rather than an estimate of the actual number of recombination-competent cRSSs across the locus.

We next estimated the frequency of selected cryptic rearrangements using fluctuation PCR, considering only sites directly interrogated with site-specific primers. Four Z43-DJβ2.1 rearrangements were observed in 2.4 µg of gDNA, corresponding to approximately 10 R/Mc for Jβ2.1 (Figure 3A). Assuming equivalent usage of all 11 Jβ gene segments, this corresponds to an estimated overall Z43 frequency of approximately 110 R/Mc (Figure 3A and Table S4). Similarly, five F-Jβ2.1 rearrangements were observed in 240 ng of DNA, corresponding to approximately 125 R/Mc for Jβ2.1 and an extrapolated frequency of approximately 1,365 R/Mc across all Jβ segments (Figure 3A and Table S4), within the same order of magnitude as Vβ30. The higher-molecular-weight product detected in this assay corresponded to direct F-Dβ2 recombination without prior Dβ-to-Jβ rearrangement (Table 1A).

**Figure 3.**
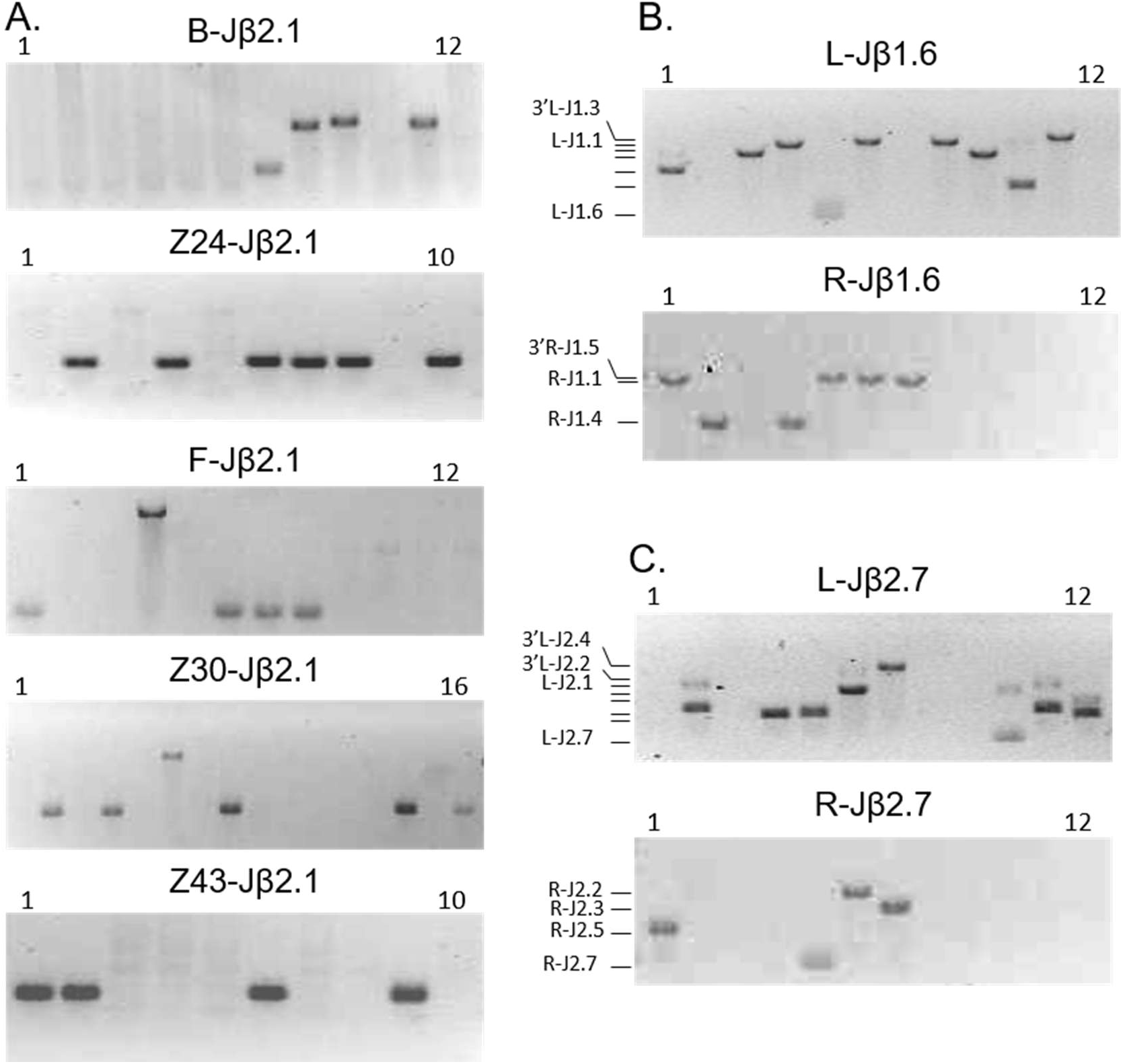
Frequencies of cryptic RSS rearrangements assessed by fluctuation PCR. (A) Representative fluctuation-PCR analyses of cryptic-site rearrangements with Jβ2.1. B–Jβ2.1 rearrangements were assessed in 12 reactions containing 480 ng of WT thymocyte gDNA each (∼960,000 cells in total); Z24–Jβ2.1 rearrangements in 10 reactions containing 240 ng each (∼400,000 cells); F–Jβ2.1 rearrangements in 12 reactions containing 20 ng each (∼40,000 cells); Z30–Jβ2.1 rearrangements in 16 reactions containing 125 ng each (∼333,333 cells); and Z43–Jβ2.1 rearrangements in 10 reactions containing 240 ng each (∼400,000 cells). (B, C) Agarose gel images showing rearrangements of the L and R cryptic sites with the Jβ1 (B) and Jβ2 (C) clusters. For each cryptic site and Jβ cluster, 12 reactions containing 500 ng of WT thymocyte gDNA each were performed, corresponding to approximately one million cells in total.

Using the same approach, recombination frequencies were estimated at approximately 165 and 198 R/Mc for Z24 and Z30, respectively; 11.5 and 34.4 R/Mc for B and 3′B; and 9 and 4.5 R/Mc for L and R, respectively (Figures 3A–C and Table S4). We also designed a forward primer specific for cV6 and tested the equivalent of one million thymocytes for recombination with Jβ2.1, but no additional cV6 rearrangements were detected, indicating a frequency below approximately 1 R/Mc.

Thus, the cryptic sites examined spanned more than three orders of magnitude in estimated recombination frequency. Their frequencies ranged from <1 R/Mc to approximately 1,365 R/Mc and were generally 1.6- to 440-fold lower than the estimated Vβ30 rearrangement frequency of 2,200 R/Mc, although site F rearranged within the same order of magnitude as Vβ30. Overall, cryptic-site frequencies were comparable between the Vβ and Vβ30–Dβ1 regions, although no site within the Vβ30–Dβ1 region reached the frequency observed for F.

For comparison with conventional Vβ rearrangement, we performed a simple extrapolation based on three observations: Vβ30 contributes approximately 0.4% of the Vβ repertoire; the cryptic sites quantified here rearranged, on average, at approximately one-tenth the frequency of Vβ30; and extrapolation of the observed site density would predict approximately 172 forward-oriented 23-cRSS-like sites across the locus. Under these assumptions, such cryptic rearrangements could theoretically correspond to approximately 7% of the number of conventional Vβ rearrangements. This calculation does not include inversional rearrangements or rearrangements involving cryptic 12-RSS-like sequences and therefore captures only one class of potential cryptic events. Consistent with the existence of additional classes, we identified three coding joints containing unconventional recombination sites on the Jβ side (Vβ10-3′Jβ2.3, 3′Z42-3′Jβ1.5 and 3′Z59-3′Jβ1.3; Table S4). Although no cRSS passing the REC threshold was identified at these positions, the coding-joint structures were consistent with RAG-mediated recombination.

This extrapolation should nevertheless be interpreted cautiously. Candidate sites were not sampled randomly, detection sensitivity varied between assays, and recombination frequencies differed substantially among individual cRSSs. The ∼7% value should therefore be regarded as illustrating the potential cumulative scale of cryptic rearrangement rather than as a quantitative estimate of its contribution to the TCRβ repertoire.

### Multiple cryptic recombination sites form a hotspot within the Bcl11b locus

To determine whether the cryptic rearrangements identified at TCRβ were restricted to antigen receptor loci, we applied the same strategy to *Bcl11b* (Figure S3A). *Bcl11b* contains two previously described cRSSs that rearrange at frequencies of approximately 23–50 R/Mc (8, 19). We detected this previously characterized rearrangement at 26.4 R/Mc (Figure S3B), consistent with the published range. Based on their REC scores, the most likely configuration was an intron 3 12-cRSS (CACAGTGATCACAGCCATTCTTACAACA; REC=0.85) paired with an intron 1 23-cRSS (CACACACATGCACACACATTACATGCCCAGCCCATATCC; REC=0.72; see (47)).

We then searched for additional forward-oriented 23-cRSSs that could undergo deletional recombination with the *Bcl11b* intron 3 12-cRSS. Twenty candidate regions located 3′ of the intron 3 cRSS were analysed by nested fluctuation PCR and designated I–XX in order of increasing distance from the intron 3 cRSS.

Of these 20 candidates, ten contained at least one predicted Z-DNA sequence (I, VII, X, XII, XIII, XV, XVI, XVII, XIX and XX), six contained a sequence with a favourable REC score upstream of the reverse primer (II, VI, VIII, XI, XIV and XVIII), and four contained both a predicted Z-DNA motif and a favourable REC score (III, IV, V and IX; Table S5). Most assays produced no detectable amplicon. The previously described intron 1 cRSS rearrangement was detected in an assay designed to interrogate a site more than 1 kb downstream of intron 1. Two additional assays, *Bcl11b* III and IV, produced products of the expected size (Figure S3C), and sequencing confirmed coding joints consistent with RAG-dependent recombination (Table 2).

**Table 2.**
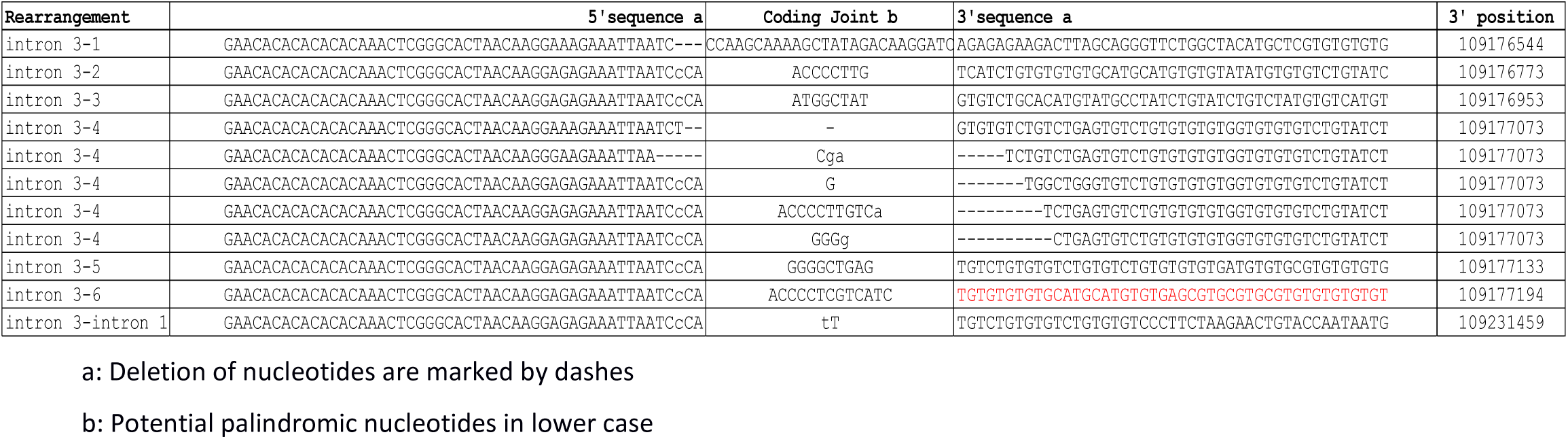
Bcl11b cryptic rearrangements coding joints.

These newly identified breakpoints clustered within a 650-bp region between chr12:109176544 and chr12:109177194 (mouse genome assembly NCBIM37). Within this interval, 11 distinct rearrangements involving six cryptic sites were detected from the equivalent of two million cells, corresponding collectively to approximately 5.5 R/Mc. The 20 assays interrogated an estimated 40 kb of sequence. Including the previously described intron 1 cRSS, the observed density was approximately 0.05 detectable 23-RSS-like sites per kb across the interrogated *Bcl11b* region, approximately five-fold lower than the density observed at TCRβ.

Thus, application of the same targeted strategy outside an antigen receptor locus identified a cluster of additional cryptic rearrangements at *Bcl11b*, demonstrating that the low-frequency RAG-mediated events detected in this study are not restricted to TCRβ.

### Local genetic and epigenetic features do not reliably predict cryptic-site recombination

Having identified 33 cryptic sites at the TCRβ locus, we next asked whether sequence or epigenetic features could distinguish sites with detectable cryptic recombination from those at which recombination was not detected. Three groups were defined (Table S3): (i) RSSs associated with functional Vβ gene segments (vRSSs); (ii) RSSs from recombining pseudogenes together with the 33 cryptic sites identified by targeted PCR and sequencing (cRSSs); and (iii) pseudogene RSSs and candidate sites at which rearrangement was not detected (nonRSSs).

As expected, REC scores clearly distinguished vRSSs from the other two groups (*P*<0.01, Wilcoxon rank-sum test; Figure 5A). However, 11 of the 33 cRSSs did not pass the REC sensitivity thresholds. For nonRSSs, the highest REC score within 300 bp downstream of the PCR primer was selected, a procedure that inherently biases this group towards higher REC scores. Despite this bias, REC score did not distinguish cRSSs from sites at which recombination was not detected, indicating that sequence-based predicted recombination potential alone was insufficient to explain low-frequency *in vivo* usage.

We further investigated whether intrinsic RSS recombination efficiency correlated with *in vivo* segment usage using the GFPi fluorescent reporter assay (48). RSSs associated with functional gene segments and pseudogenes were individually paired with the Dβ2 12-RSS (Supplementary Text, Table S6 and Figure S4). Notably, the Vβ8, Vβ25 and Vβ28 RSSs exhibited greater *in vitro* recombination efficiency than the Vβ30 RSS, despite the absence of detectable *in vivo* rearrangement in our assays. Similarly, neither the presence, score nor density of predicted Z-DNA motifs was associated with detectable cryptic recombination, with Z-DNA motifs occurring in both cRSS and nonRSS groups (Table S3). Together, these results show that neither predicted RSS quality nor the intrinsic recombination efficiency measured in the reporter assay was sufficient to predict low-frequency *in vivo* recombination.

We next examined whether the local epigenetic environment could distinguish cRSSs from nonRSSs. Publicly available datasets from double-negative thymocytes were analysed for H3K4me1, H3K4me2, H3K4me3, H3K27me3, H3K36me3, H3K27ac, H3K9me2, H3ac, FAIRE, CTCF, Pol II, P300, and sense and antisense germline transcription (27, 35, 37, 38). We additionally measured interactions between the TCRβ enhancer Eβ and the remainder of the locus by 4C (see Methods). None of the individual features examined reliably distinguished cRSSs from sites at which rearrangement was not detected (Figure 4 and Figure S5). Indeed, many cryptic sites showed low levels of chromatin-activation marks, whereas some sites without detectable recombination showed comparatively high levels (examples in Figure S6). In contrast, several of these features more clearly distinguished functional vRSSs from the cRSS and nonRSS groups, consistent with the strong correlations among many of the epigenetic variables examined (Figure S7).

**Figure 4.**
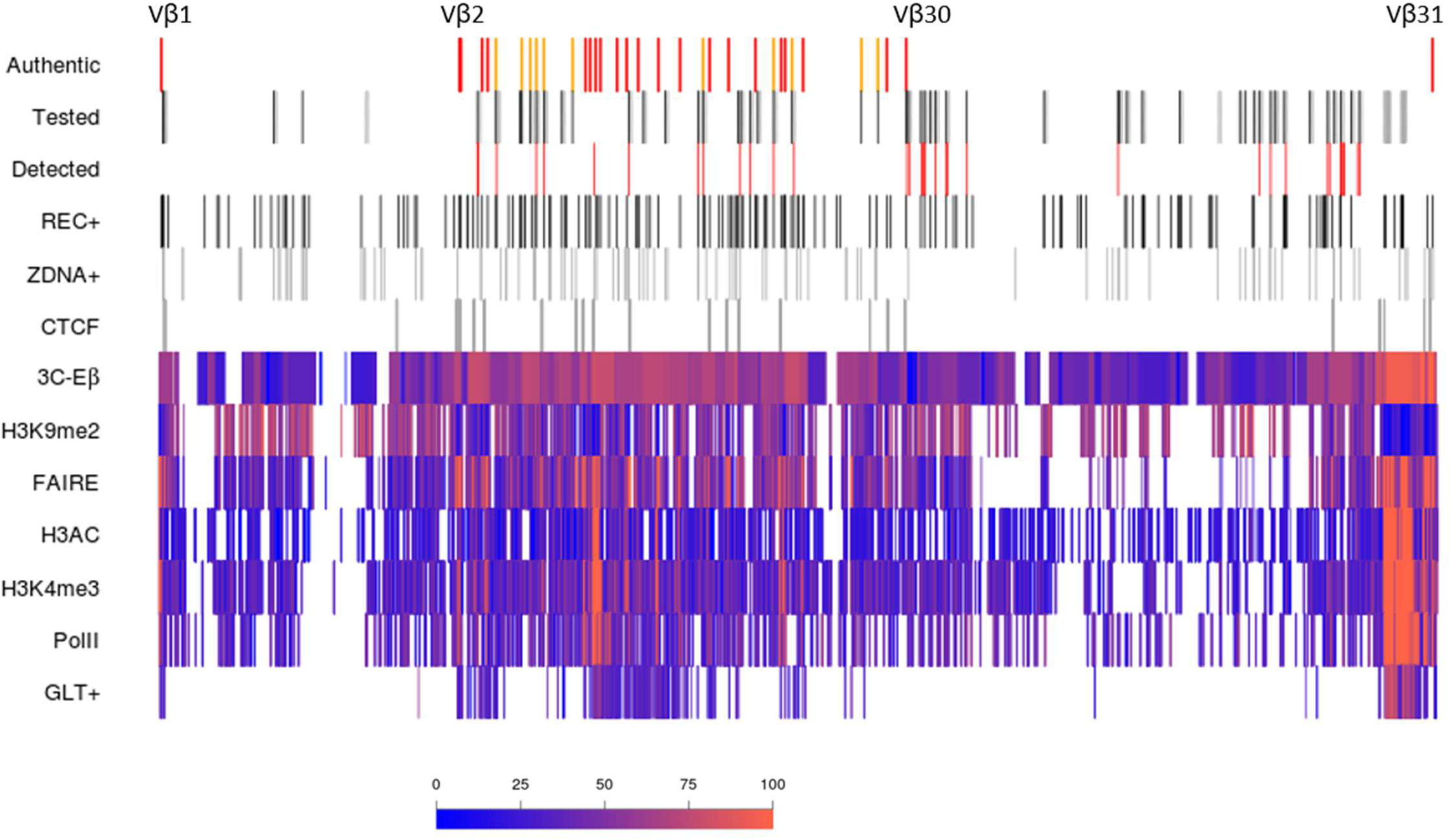
Cryptic RAG-mediated rearrangements across the TCRβ locus and their genetic and epigenetic context. Functional Vβ gene segments are shown in red and Vβ pseudogenes in orange. Vβ1 and Vβ31 are located at the extreme 5′ and 3′ ends of the locus, respectively, whereas the remaining Vβ segments are clustered within a ∼230-kb region. Regions interrogated for cryptic rearrangements are shown in grey, extending 2 kb downstream of each designed primer, with black lines indicating the specific candidate cRSSs targeted. Cryptic recombination events detected by PCR and confirmed by sequencing are shown as red lines. REC+ indicates sites passing the stringent REC threshold (putative cRSSs), and Z-DNA+ indicates regions computationally predicted to have Z-DNA-forming potential. CTCF indicates peaks of ChIP-seq signal enrichment in double-negative thymocytes. The remaining tracks show the relative signal intensity of the indicated epigenetic features across the locus in double-negative thymocytes; white regions indicate absence of available data. GLT+ indicates positive-strand germline transcription, measured by stranded total RNA sequencing.

We therefore integrated these genetic and epigenetic variables using Random Forest classification. Across 100 repeated models, OOB predictions correctly classified >90% of vRSSs, whereas error rates remained >30% for discrimination of cRSSs and nonRSSs (Figure 5). Several of the 37 sites in the cRSS group (33 cryptic sites and four rearranging pseudogene RSSs) were misclassified as either vRSSs or nonRSSs, whereas misclassified nonRSSs were only rarely assigned to the vRSS group. Thus, the features examined contained sufficient information to distinguish canonical functional RSSs relatively well, but were insufficient to reliably separate low-frequency cryptic substrates from candidate sites at which rearrangement was not detected. A similar pattern was observed at *Bcl11b*. The newly identified recombination hotspot showed low levels of the activating chromatin marks H3ac and H3K4me3 (Figure 6), indicating that strong enrichment for these marks at both recombination partners is not required for detectable cryptic recombination at the population level. The hotspot nevertheless contained the highest-scoring predicted Z-DNA sequence within the gene (Z-score 301 over 92 nucleotides) and the highest local density of Z-DNA motifs, with 245 nucleotides predicted to adopt Z-DNA within the 674-nucleotide interval encompassing all breakpoints. Conversely, several *Bcl11b* candidate sites at which rearrangement was not detected displayed comparatively high levels of activating chromatin marks.

**Figure 5.**
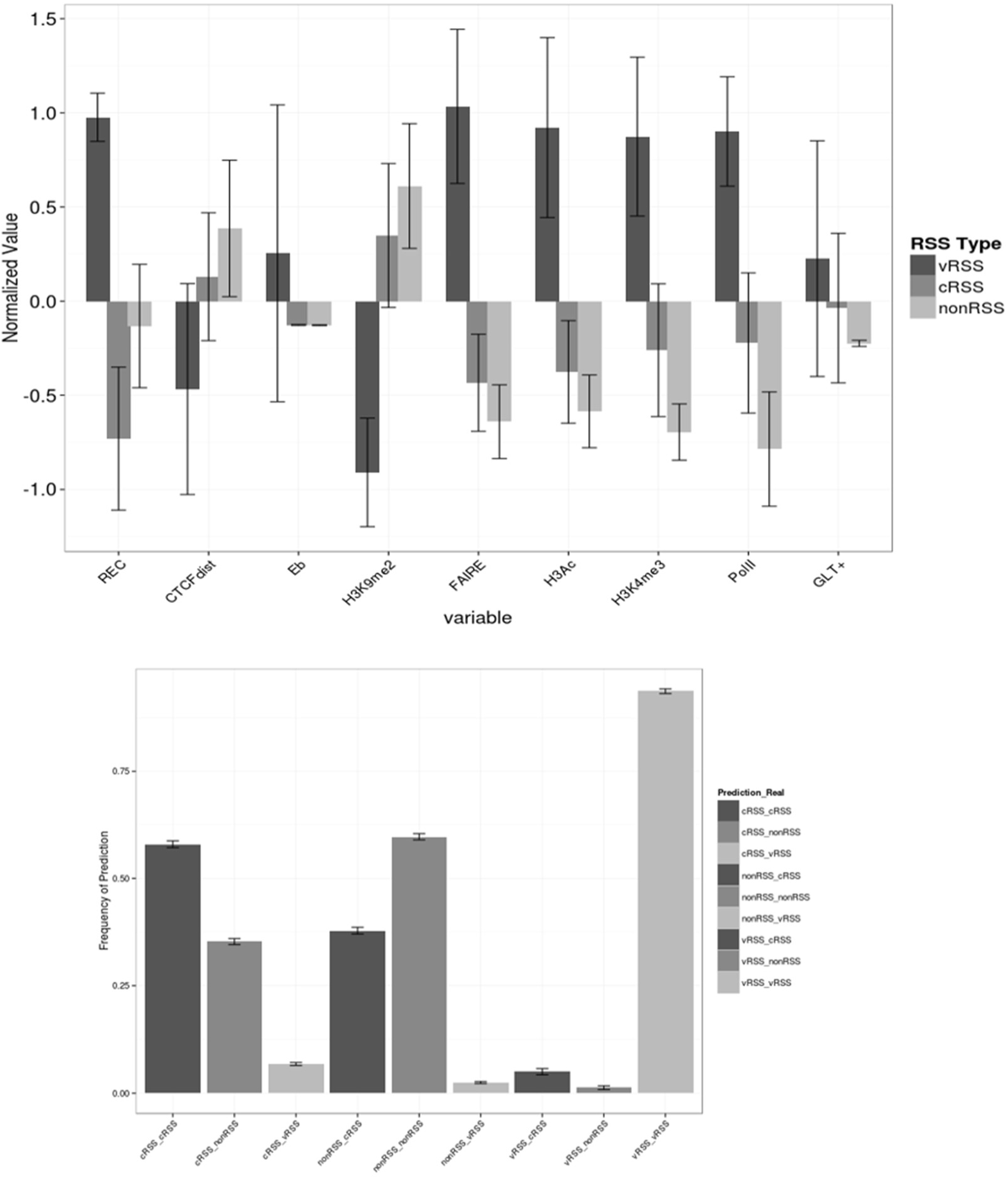
Genetic and epigenetic features of recombination sites and their ability to predict cryptic recombination. (A) Genetic and epigenetic features of genomic sites grouped according to their recombination status. Features include REC score, distance (bp) to the nearest CTCF peak, and signal intensity for the indicated epigenetic marks (see Methods). All variables were transformed to standard scores. Error bars indicate 95% confidence intervals of the mean. (B) Classification performance of Random Forest models. Bars show the mean frequency of out-of-bag (OOB) classification outcomes across 100 repeated Random Forest models, grouped according to observed and predicted classes. Error bars indicate 95% confidence intervals across the 100 repetitions and therefore represent run-to-run variability of the classifier on the fixed dataset rather than sampling variability of the underlying population.

**Figure 6.**
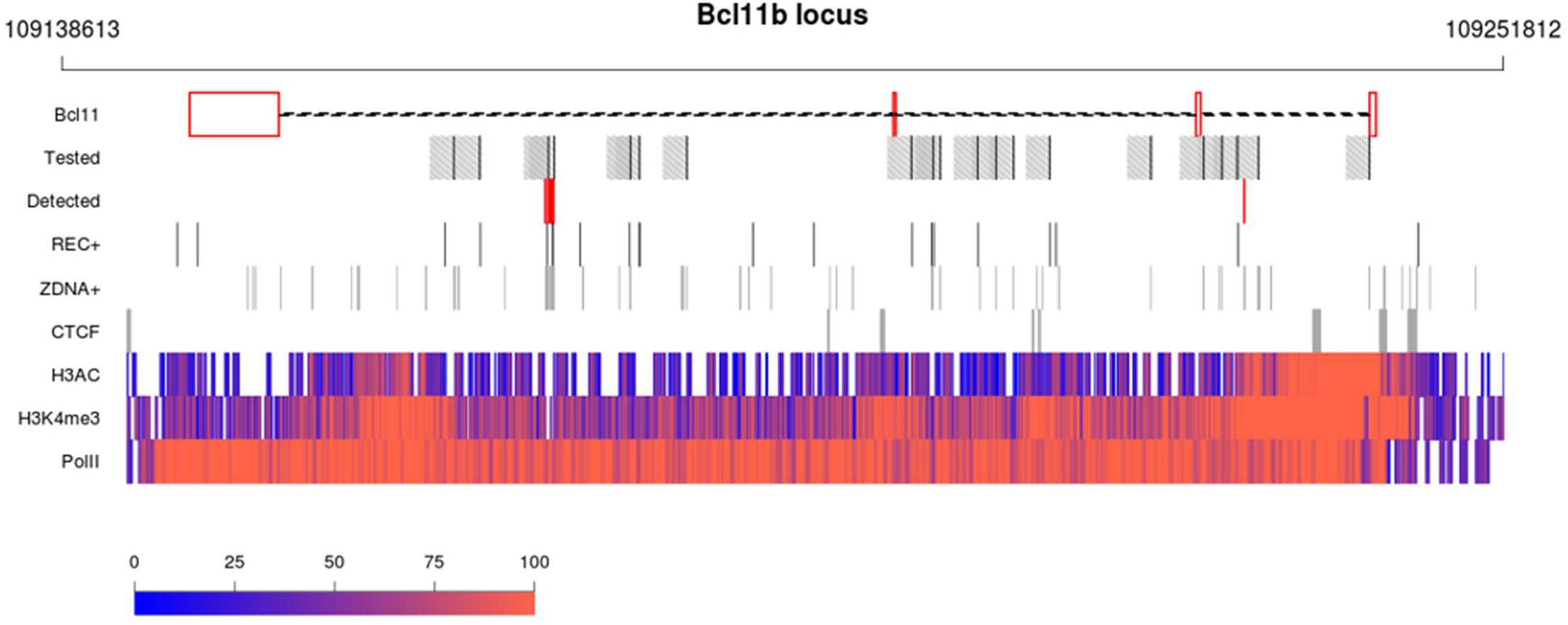
Cryptic RAG-mediated rearrangements across the Bcl11b locus and their genetic and epigenetic context. Bcl11b exons are shown as red boxes, with introns represented by dashed connecting lines; note that *Bcl11b* is located on the reverse strand. Regions interrogated for cryptic rearrangements are shown in grey, extending 2 kb downstream of each designed primer, with black lines indicating the specific candidate cRSSs targeted. Cryptic recombination events detected by PCR and confirmed by sequencing are shown as red lines. REC+ indicates sites passing the stringent REC threshold (putative cRSSs), and Z-DNA+ indicates regions computationally predicted to have Z-DNA-forming potential. CTCF indicates peaks of ChIP-seq signal enrichment in double-negative thymocytes. The remaining tracks show the relative signal intensity of the indicated epigenetic features across the locus in double-negative thymocytes; white regions indicate absence of available data. Unlike Figure 4, 3C, FAIRE and H3K9me2 are not shown because data were unavailable for this locus. The scale is identical to that used in Figure 4.

Overall, these analyses indicate that local RSS sequence, predicted Z-DNA formation and population-level epigenetic accessibility each capture aspects of RAG substrate biology, but none – alone or in combination in the models tested here – reliably predicts which cryptic sites undergo detectable low-frequency rearrangement *in vivo*.

## DISCUSSION

While investigating the recombination potential of Vβ pseudogenes, we identified numerous previously uncharacterized cryptic RAG-mediated rearrangements at the TCRβ locus. In total, 33 cryptic sites were detected, distributed across both the predominantly accessible Vβ region and the comparatively inaccessible Vβ30–Dβ1 intergenic region. Within the ∼106 kb effectively interrogated, 26 cryptic sites were identified, corresponding to an observed density of approximately 0.25 cryptic 23-RSS-like sites per kb. Recombination frequencies varied substantially between sites and were generally lower than that of the functional Vβ30 segment, although some sites recombined at a similar order of magnitude. We further identified additional cryptic rearrangements at the Bcl11b locus, indicating that such events are not restricted to antigen receptor loci. Together, these observations reveal a broad background of low-frequency RAG-mediated DNA rearrangement during normal T-cell development.

Our approach was designed to detect completed rare rearrangements rather than RAG cleavage itself and therefore provides information complementary to subsequent genome-wide approaches. High-throughput analyses have since demonstrated extensive RAG off-target activity within antigen receptor loci and directly detected RAG-dependent DNA breaks at cryptic sites in developing lymphocytes (20, 21). In particular, RAG off-target rearrangements have been identified within the Tcrd locus in developing T cells, while END-seq detected cryptic RAG-dependent breaks in primary thymocytes. These later observations support the conclusion that low-frequency off-target RAG activity is a genuine feature of normal lymphocyte development rather than being restricted to transformed or malignant cells.

Several limitations of our experimental strategy should nevertheless be considered. The abundance of canonical Vβ rearrangements in total thymocytes generated large numbers of authentic V(D)J products despite the use of (semi-)nested PCR, reducing sensitivity for rare cryptic events. Repetitive and duplicated sequences within the TCRβ locus also constrained primer design and, in some cases, prevented unambiguous assignment of rearrangement sites. In addition, the Nextera-based sequencing strategy may have under-represented short PCR fragments. Candidate sites were selected on the basis of predicted RSS or Z-DNA features rather than sampled randomly across the locus, and detection sensitivity depended on primer position and amplicon length. Consequently, the density and frequencies reported here should not be interpreted as quantitative estimates of cryptic recombination across the entire TCRβ locus. Rather, they demonstrate that cryptic rearrangements are readily detectable at multiple sites when specifically interrogated and suggest that their true number is likely to be greater than captured by this targeted approach.

A central question was whether cryptic target selection could be predicted from local sequence or chromatin features. Neither REC score nor predicted Z-DNA formation reliably distinguished sites with detectable rearrangement from sites at which rearrangement was not detected. Similarly, integration of multiple population-level epigenetic features and chromosomal interactions failed to provide reliable discrimination between these groups. Notably, cryptic rearrangements were detected within the Vβ30–Dβ1 region despite its predominantly inaccessible chromatin profile in DN thymocytes. These observations indicate that local RSS resemblance and steady-state chromatin accessibility, although important determinants of physiological V(D)J recombination, are insufficient on their own to explain low-frequency cryptic target selection.

Subsequent studies provide a mechanistic framework that may help explain this observation. RAG bound at recombination centres can scan surrounding chromatin and encounter potential substrates in an orientation-dependent manner within chromosomal loop domains (20). CTCF-binding elements can modulate the accessibility and interaction of potential RAG substrates with recombination centres (28), and cohesin-mediated loop extrusion has subsequently emerged as a fundamental mechanism presenting chromatin to recombination-centre-bound RAG (29). Importantly, experimentally altering locus orientation or loop-extrusion constraints changes the utilization not only of canonical RSSs but also of cryptic RSSs (30). Thus, cryptic target selection is likely to reflect an interaction between RSS sequence and orientation, local chromatin state, recombination-centre activity and higher-order chromatin architecture rather than any single local feature. This framework also changes the interpretation of our inability to relate cryptic recombination simply to genomic distance. Previous work had suggested that proximity to the recombination centre influences RSS usage (26, 27, 49, 50). In our dataset, neither linear distance from the cryptic site nor distance to the nearest CTCF site correlated with detectable rearrangement. Five sites positioned between 12 kb upstream and 9.6 kb downstream of a proposed anchor for distal Vβ interactions with the DJ recombination centre had similar epigenetic profiles and REC scores, yet only two underwent detectable rearrangement. Rather than arguing that locus organization is unimportant, subsequent evidence suggests that simple linear distance or proximity to an individual CTCF site is unlikely to capture the dynamic contacts generated by chromatin scanning and loop extrusion. Our findings are therefore consistent with a model in which the probability of cryptic recombination depends on whether a potential substrate is productively presented to RAG within the appropriate three-dimensional and orientation-dependent context.

Population-level epigenetic measurements may likewise obscure transient or cell-specific states relevant to rare events. RAG2 recognition of H3K4me3 provides an important link between chromatin accessibility and recombination (22), yet high H3K4me3 enrichment was not a prerequisite for the cryptic rearrangements detected here, consistent with previous observations that transcriptional or chromatin activity alone does not determine susceptibility to RAG-mediated rearrangement (51). One possibility is that rare cryptic events arise from transiently permissive chromatin configurations that are poorly represented in bulk epigenomic datasets, consistent with the proposed contribution of stochastic epigenetic variation to antigen receptor gene rearrangement (69). However, our data do not directly demonstrate such stochastic states, and the subsequent discovery of chromatin scanning and loop extrusion provides an alternative, non-mutually exclusive explanation. Resolving the relative contribution of transient local chromatin states and dynamic locus architecture will require approaches capable of linking RAG cleavage or rearrangement to chromatin state at single-cell or single-allele resolution.

Cryptic RSSs within antigen receptor loci are not necessarily deleterious. Defined cRSSs participate in processes such as VH replacement (52), diversification of the primary VH repertoire (53), and Igκ locus inactivation (54–56), while additional conserved cryptic sites have been described whose functions remain less clear (57, 58). The sites identified here differ from these conserved functional cRSSs: they show little apparent conservation in sequence or position relative to gene segments and, in the orientation examined, cannot mediate conventional gene-segment replacement. Their rearrangement frequencies nevertheless overlap those reported for cryptic sites at non-antigen receptor loci, including HPRT, Notch1 and Bcl11b (6–8, 19). Moreover, because our analysis focused primarily on forward-oriented 23-RSS-like sequences, additional classes of cryptic events-including reverse-oriented sites and 12-RSS-like substrates-were not systematically interrogated. The biological consequences of most such rare rearrangements remain uncertain. Potentially deleterious RAG-mediated lesions are constrained by DNA-damage surveillance mechanisms, including ATM-dependent responses (59), while events without a selective consequence are intrinsically difficult to detect unless specifically assayed. Importantly, genome-wide studies have since shown that although RAG associates with many regions of the lymphocyte genome, productive off-target cleavage is substantially more restricted (47). Thus, the presence of potential cRSSs alone does not imply widespread genomic damage. Rather, our findings, together with subsequent genome-wide studies, support a model in which RAG generates a background of low-frequency off-target rearrangements whose occurrence is constrained by multiple layers of sequence recognition, chromatin accessibility, locus architecture and DNA-damage control.

In conclusion, our study identifies a diverse set of completed cryptic RAG-mediated rearrangements in normal developing thymocytes and demonstrates that local RSS sequence and population-level epigenetic features are insufficient to predict which potential sites undergo detectable recombination. Subsequent discoveries of orientation-dependent RAG scanning and cohesin-mediated loop extrusion provide a mechanistic context for these findings and emphasize the importance of higher-order chromatin organization in determining RAG target choice. Together, these observations illustrate how the mechanisms required to generate antigen receptor diversity inevitably create opportunities for low-frequency off-target DNA rearrangement, while highlighting the multiple layers of regulation that normally preserve lymphocyte genome integrity.

## MATERIAL AND METHODS

### Mice

Wild-type (WT) and RAG2-deficient (RAG2-KO) mice were used in this study. Mice were 6–10 weeks old at the time of sacrifice, after which thymi were collected. All experiments were conducted in accordance with guidelines for the care and use of animals under a protocol approved by the Instituto Gulbenkian de Ciência Animal Care and Use Committee.

### PCR assays and Southern blotting

For dilution-series PCR, five-fold serial dilutions of genomic DNA (gDNA) from WT total thymocytes and early DN3 (DN3E) thymocytes were amplified, using input amounts ranging from 100 ng to 0.8 ng.

To analyse pseudogene usage, semi-nested fluctuation PCR was performed by repeating the amplification of each pseudogene with both DJβ clusters 11–22 times, using 60–120 ng of gDNA from WT or RAG2-KO total thymocytes per reaction. A total of 2,640 ng of gDNA from each genotype was tested for each pseudogene. Amplicons were subsequently analysed by Southern blotting using ^32P-labelled Jβ-specific probes, as previously described (60). Vβ30–Jβ rearrangements were amplified using the same approach as a control. In addition, rearrangements involving all Jβ1 and Jβ2 gene segments were amplified using 6 ng of gDNA per reaction, and rearrangements of Vβ30 with the first gene segment of each cluster (Jβ1.1 and Jβ2.1) were amplified using 20 ng of gDNA per reaction. To estimate the recombination frequencies of Vβ10 and Vβ22, 22 and 11 PCR reactions, respectively, were performed using 6 ng of gDNA per reaction.

For cRSS analysis within the Vβ region, a total of 6 µg of WT total-thymocyte gDNA was tested for each REC candidate (A– G) by semi-nested fluctuation PCR. Sixteen reactions containing 375 ng of gDNA each were used to amplify rearrangements with Jβ1.1 or Jβ2.1 (16). Additional amplifications of rearrangements involving the complete clusters, using the reverse nested primers Jβ1.6 and Jβ2.7, were performed using 720 ng of gDNA for each cluster. Amplicons were subsequently analysed by Southern blotting. The eight Z-motif candidates were tested by semi-nested fluctuation PCR using 240 ng of WT thymocyte gDNA per reaction, for a total of 2.4 µg, to detect rearrangements with Jβ2.1. Additional amplifications of Z30– Jβ1.6 and Z30–Jβ2.7 rearrangements were performed using 6 ng of WT thymocyte gDNA per reaction, for a total of 72 ng. The 26 sites within the Vβ30–Dβ1 region were tested by nested PCR with Jβ1.6 and Jβ2.7. Twelve reactions containing 500 ng of WT total-thymocyte gDNA per reaction were performed.

For each PCR reaction, DNA was amplified using 1 U of Flexi GoTaq polymerase (Promega) in 1× GoTaq buffer containing 2 mM MgCl₂, 0.2 mM dNTPs and 0.2 mM of each primer. PCR conditions were 2 min at 94°C, followed by 30 cycles of 30 s at 94°C, 30 s at 59°C and 1 min at 72°C, with a final extension of 10 min at 72°C. One microlitre of the first PCR product was subjected to a further 30 cycles using either the same forward primer (semi-nested PCR) or an internal forward primer (nested PCR), together with an internal Jβ1.1, Jβ2.1, Jβ1.6 or Jβ2.7 primer. Pseudogene-specific forward primers and the first set of Jβ1.1 and Jβ2.1 primers were those previously described (27). All other primers and probes, including an additional set of primers for the Vβ6, Vβ10, Vβ18, Vβ22 and Vβ25 pseudogenes (Table S1A), were designed using Primer3 (61). Half of each PCR product was analysed by agarose gel electrophoresis. gDNA from RAG2-KO thymocytes was used as a negative control.

At the Bcl11b locus, the previously reported deletion was amplified by nested PCR using the oligonucleotides and PCR conditions described previously (19). Twenty pairs of reverse primers, corresponding to the first and second PCR reactions, were additionally designed using Primer3 (61) (Table S1A) and tested in combination with the forward primers described by Sakata et al. (19). Twelve reactions containing 500 ng of WT total-thymocyte gDNA per reaction were performed. PCR conditions were as described above.

### Deep sequencing of pooled PCR products

Half of the PCR products from reactions exhibiting at least one positive band by agarose gel electrophoresis were pooled and sequenced at the IGC sequencing facility using an Illumina MiSeq instrument and a standard Nextera protocol, generating 300-bp paired-end reads (∼1 million reads per sample).

For each sample, low-quality bases (p<99%) were trimmed from read ends using seqtk. Overlapping paired-end reads were merged, where possible, to generate longer contigs using FLASH (62). High-quality contigs were aligned to the reference sequence using LASTZ (63). Alignments were retained when they showed at least 85% sequence identity to the reference and consisted of two non-overlapping sub-alignments of at least 50 bp each, together covering at least 90% of the contig. Alignments with similar split positions and similar putative nucleotide additions or deletions at the non-homologous end-joining (NHEJ) junction, allowing up to three differences, were then clustered.

Raw sequencing data are available from NCBI under accession PRJNA400571.

### DNA sequence analysis

The TCRβ locus and Bcl11b gene were scanned using ABCC to identify sequences containing predicted Z-DNA motifs and their associated Z-scores (64).

The REC algorithm was used to calculate REC scores for candidate cryptic sites and for sequences downstream of breakpoints identified by PCR and deep sequencing. For each tested potential RSS, the most probable cRSS was identified. For sites with detectable recombination, the first cRSS downstream of the breakpoint that passed the REC sensitivity threshold was selected. For sites at which recombination was not detected, the sequence with the highest REC score within 300 bp downstream of the primer was selected.

To calculate Z-density scores, the locus was divided into 500-bp windows with a 100-bp step. For each window, all overlapping Z-DNA motif regions, together with their scores and lengths, were identified. To account for partial overlap of a Z-DNA region with a given window, the Z-score per base pair for each motif was multiplied by the number of overlapping base pairs. The Z-density score for each window was calculated as the sum of these values divided by the window length.

### Circularized chromosome conformation capture (4C)

4C-seq experiments were performed as previously described (65). Libraries were prepared from RAG2-deficient (RAG2-KO) thymocytes using the NlaIII/DpnII restriction enzyme combination, with Eβ as the viewpoint. Primer sequences are provided in Table S1B. Two replicate experiments were sequenced and merged for downstream analysis.

4C-seq data were processed as previously described (66). For quantification, normalized reads-per-million (RPM) data were smoothed using a running-mean approach. The 40%, 50% and 60% quantiles were subsequently smoothed and interpolated using the R loess function implemented in Basic4Cseq (67).

### Epigenetic data analysis

Processed ChIP-chip data from double-negative (DN) thymocytes for H3K27me3, H3K36me3, H3K4me1, H3K4me2, H3K4me3 and H3K9me2, together with RNA polymerase II (Pol II) ChIP-seq data, were obtained from (35). Processed P300 and H3K27ac ChIP-chip data and H3ac ChIP-seq data were obtained from GEO under accession numbers GSE49234 and GSE48817 (27). Germline transcription and FAIRE data were obtained from references (68) and (27) respectively. Processed datasets were used as provided because they had already been normalized and log-transformed. For the 3C, Pol II, H3ac, H3K4me1 and H3K4me3 datasets, the logarithm of the signal was used.

For each site of interest (functional Vβ RSSs and tested cRSSs), a 1,000-bp region centred on the site was generated. Each region was divided into 50-bp bins, and the mean signal within each bin was calculated to generate signal profiles across the region. Following manual inspection of the distribution of profiles across loci, a window was selected for each epigenetic feature. For FAIRE, the mean signal within the 250 bp immediately 3′ of the RSS was used; for H3ac, the 500 bp immediately 5′ of the RSS was used; and for all remaining features, the entire 1,000-bp window was used. The mean signal for each variable at each locus was subsequently used for correlation analyses and Random Forest classification. Random Forest classification was performed in R using the randomForest package (69). Loci were assigned to one of three classes (vRSS, cRSS or nonRSS), and all remaining variables were used as predictors. Forests were grown using the default settings of 500 trees and √p variables sampled at each split, where p is the number of predictors. Classification performance was assessed using out-of-bag (OOB) predictions. For each forest, the OOB confusion matrix was normalized by row to obtain, for each observed class, the proportion of sites assigned to each predicted class. The analysis was repeated 100 times on the same dataset (random-number generator seed 111), with repetitions differing only in the internal randomization of the Random Forest algorithm. Classification frequencies were summarized across the 100 repetitions as means with 95% confidence intervals. These intervals represent run-to-run variability of the classifier on the fixed dataset rather than sampling variability of the underlying population.

### Gene-segment homology analysis

For each candidate cRSS, the 295 nucleotides immediately upstream of the site, corresponding to the average length of functional Vβ gene segments, were queried against the Mouse Ensembl Ab-initio Proteins database (version 73) using BLASTX optimized for distant homologies. The nucleotide sequences, including introns, of three functional gene segments (Vβ2, Vβ3 and Vβ4) and three pseudogene segments (Vβ7, Vβ8 and Vβ9) were used as positive controls. A tested sequence was considered a potential gene segment if it produced a hit to any other annotated gene segment with an E-value <1 × 10^−5.

### Fluorescence-based in vitro recombination assay

The CFP-GFPi reporter construct was described previously in Trancoso et al. (1). Briefly, the 12-RSS (either Dβ2 or ConS12) was cloned into the BX site, whereas each Vβ 23-RSS was cloned into the HE site. The 12-RSS was also cloned without a 23-RSS partner to determine background GFP expression independently of the Vβ RSS.

HEK293T cells were transfected as previously described (1) using Lipofectamine (Invitrogen). Briefly, 60,000 cells were plated in 24-well plates 24 h before transfection. Cells were transfected with 1 µg of the GFPi-CFP construct in the presence or absence of a mixture containing 0.4 µg of RAG1 and 0.35 µg of RAG2 plasmids. Medium was replaced 16 h after transfection. At 62 h post-transfection, cells were stained with propidium iodide (Fluka) and acquired on an LSRFortessa flow cytometer (BD Biosciences).

Data were analysed using FlowJo (Tree Star). Analysis was restricted to propidium iodide-negative 293T singlets within the highest 15% of CFP-expressing cells. For each construct, the GFP^+ gate was positioned to include 2% of cells in the RAG-negative sample and then applied unchanged to the corresponding RAG-positive sample within the 15% CFP gate. For each RSS pair, the recombination efficiency (RE) score was calculated by subtracting the percentage of GFP^+ cells in the RAG-negative sample (∼2%) from the percentage of GFP^+ cells in the corresponding RAG-positive sample.

## Acknowledgments

We wish to thank Thiago Carvalho, Jorge Carneiro, Vasco M. Barreto, Alekos Athanasiadis and members of our laboratories for helpful discussions. The authors acknowledge funding by the Instituto Gulbenkian de Ciencia and by the Fundacao para a Ciencia e Tecnologia grant PTDC/BIA-GEN/116830/2010. MBDP was funded by a postdoctoral FCT fellowship SFRH/BPD/65292/2009. The authors declare they do not have any competing interests.

## SUPPLEMENTARY MATERIAL

### Supplementary materials and methods

#### fluorescence-based *in vitro* recombination assay

The CFP-GFPi reporter construct used in this study was described previously (48). Briefly, the 12-RSS (either Dβ2 or ConS12) was cloned into the BX site, whereas each Vβ 23-RSS was cloned into the HE site. The 12-RSS was also cloned without a 23-RSS partner to determine the background level of GFP expression independently of the Vβ RSS.

HEK293T cells were transfected as previously described using Lipofectamine (Invitrogen). Briefly, 60,000 cells were plated in 24-well plates 24 h before transfection. Cells were transfected with 1 µg of the GFPi-CFP construct in the presence or absence of a mixture containing 0.4 µg of RAG1 and 0.35 µg of RAG2 plasmids. The medium was replaced 16 h after transfection. At 62 h post-transfection, cells were stained with propidium iodide (PI; Fluka) and acquired on an LSRFortessa flow cytometer (BD Biosciences).

Data were analysed using FlowJo (Tree Star). Analysis was restricted to PI-negative 293T singlets within the highest 15% of CFP-expressing cells. For each construct, the GFP-positive gate was positioned to include 2% of cells in the RAG-negative sample and was then applied unchanged to the corresponding RAG-positive sample within the 15% CFP gate. For each RSS pair, the recombination efficiency (RE) score was calculated by subtracting the percentage of GFP-positive cells in the RAG-negative sample (∼2%) from the percentage of GFP-positive cells in the corresponding RAG-positive sample.

### Supplementary results

#### assignment of an ambiguously amplified cryptic rearrangement to Z56

One set of coding-joint (CJ) sequences could potentially have originated from PCR products generated using either of two primer sets, Z56 or T, because the sequences aligned downstream of both primers within the TCRβ locus. The Z56 primer was located 91 bp 5′ of the CJ, and the potential cRSS downstream of this primer passed the REC sensitivity threshold. In addition, agarose gel electrophoresis of the Z56 PCR products revealed bands of several sizes consistent with those of the rearrangements subsequently identified by sequencing. In contrast, the T primer was located 323 bp 5′ of the CJ, the corresponding potential cRSS did not pass the REC sensitivity threshold, and no bands were detected by agarose gel electrophoresis. Together, these observations support assignment of these rearrangements to the Z56 cryptic site.

### Supplementary discussion

#### cryptic rearrangements are consistent with constraints beyond the 12/23 rule

Our results suggest that cryptic recombination sites, including those associated with predicted Z-DNA motifs, may be subject to constraints similar to those operating at functional RSSs beyond the classical 12/23 rule. To investigate this, we searched coding-joint sequences for stretches of at least five consecutive nucleotides matching either the Dβ1 or Dβ2 segments.

Sequences matching Dβ segments were identified in the majority of coding joints involving pseudogenes (70%; 49 of 70 sequences; Table S2) and cRSSs (69%; 174 of 252 sequences; Table S4). These proportions were comparable to that observed among rearrangements involving the functional Vβ3 gene segment (66.7%; 307 of 460 sequences; data not shown). Thus, cryptic and pseudogene-associated rearrangements showed a similar prevalence of Dβ sequence contribution to that observed for a functional Vβ segment.

We also identified three rearrangements in which the cryptic site was joined directly to a Dβ segment before Dβ-to-Jβ joining (1.2%; 3 of 252 coding joints; Table 1). Although infrequent, the presence of these intermediates demonstrates that cryptic sites can engage Dβ segments before completion of Dβ-to-Jβ rearrangement and is consistent with previous evidence that D-to-J recombination is not invariably completed before V-to-DJ recombination (70).

### SUPPLEMENTARY FIGURES

**Figure S1.**
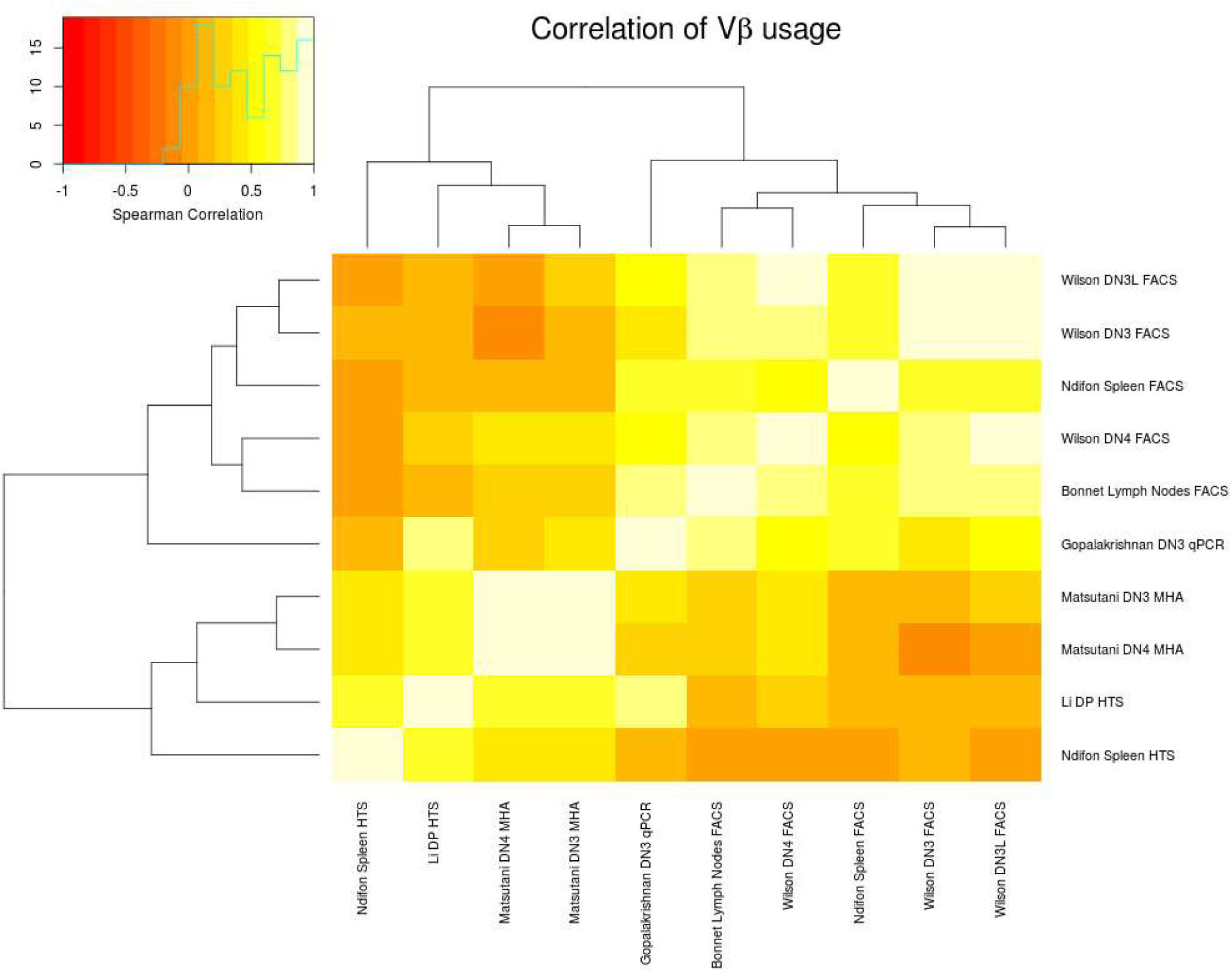
Correlations between TCR-Vβ repertoires across studies, developmental stages and analytical methods. Pairwise correlations between published TCR-Vβ repertoires are shown. Within individual studies, repertoires measured at different developmental stages were highly correlated (Wilson *et al.*; Matsutani *et al.*). In contrast, repertoires measured using different analytical approaches showed lower concordance, as illustrated by comparison of the HTS- and FACS-derived repertoires reported by Ndifon *et al.* Repertoires generated using the same analytical approach across independent studies showed greater similarity, as illustrated by the FACS-based datasets from Ndifon *et al.* and Bonnet *et al*.

**Figure S2.**
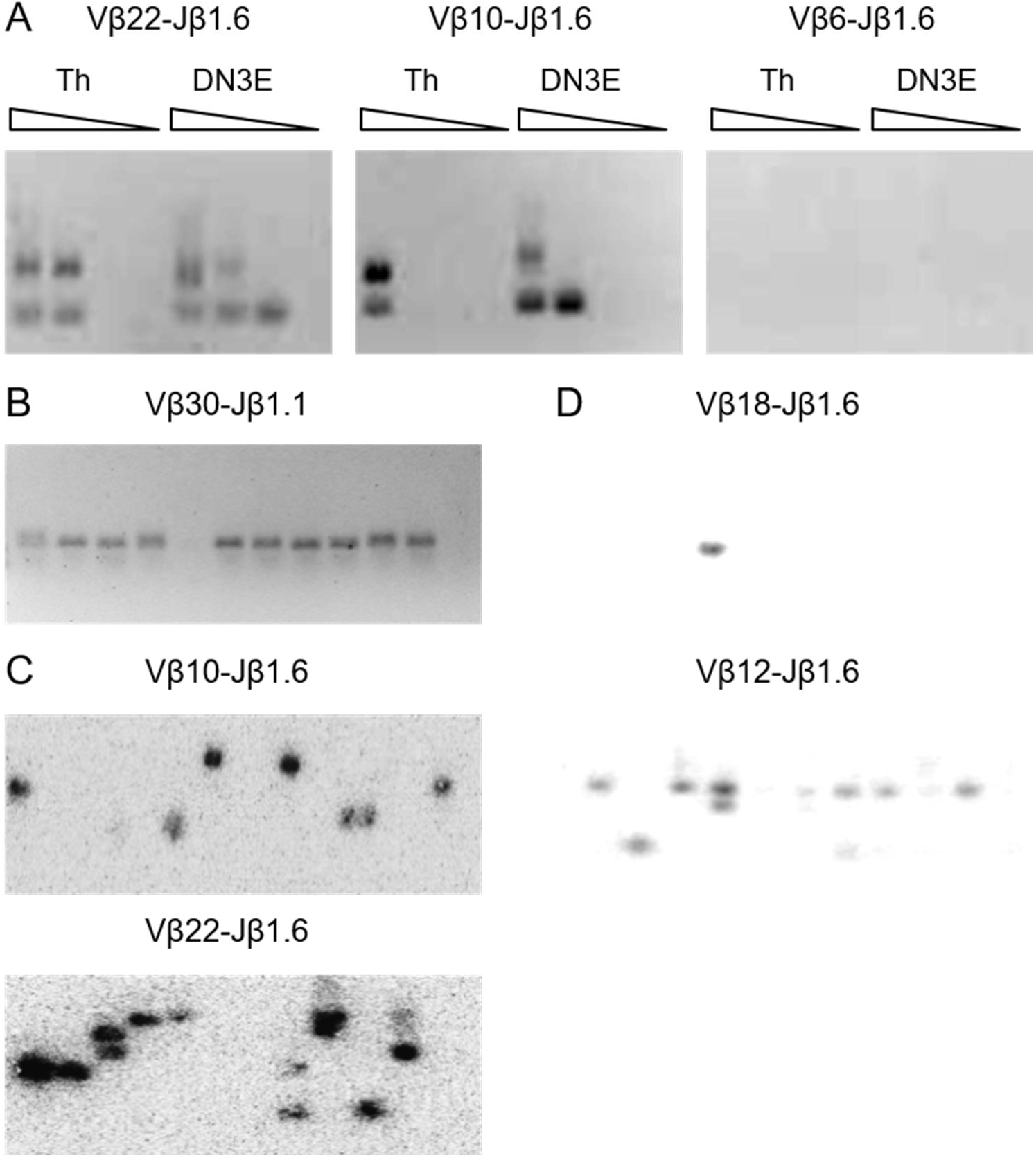
Detection of low-frequency Vβ pseudogene rearrangements by semi-nested PCR. (A) Agarose gel images of PCR amplification of five-fold serial dilutions of WT thymocyte gDNA, from 100 ng to 0.8 ng per reaction, showing Vβ22, Vβ10 and Vβ6 rearrangements with Jβ1.6. (B) Agarose gel image of 12 PCR reactions amplifying Vβ30–Jβ1.1 rearrangements, using 20 ng of gDNA per reaction. (C) Phosphorimager analysis of Southern blots from 23 and 11 PCR reactions amplifying Vβ10–Jβ1.6 and Vβ22–Jβ1.6 rearrangements, respectively, using 6 ng of gDNA per reaction. (D) Autoradiographs of Southern blots from 11 PCR reactions amplifying Vβ18–Jβ1.6 and Vβ12–Jβ1.6 rearrangements, using 60 ng of WT thymocyte gDNA per reaction.

**Figure S3.**
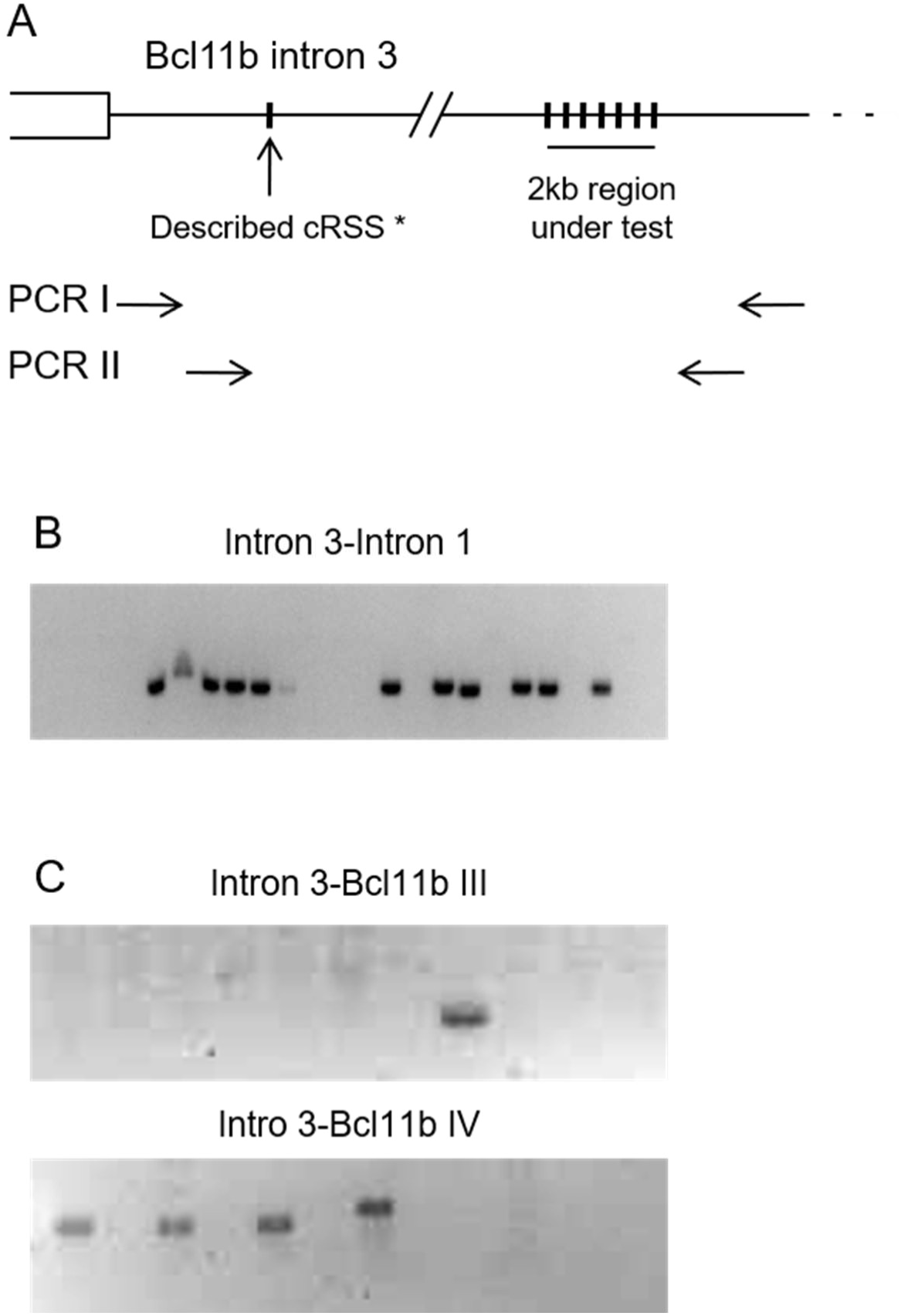
Detection and quantification of cryptic RAG-mediated rearrangements at the Bcl11b locus. (A) Schematic representation of the nested-PCR strategy used to detect cryptic RSS rearrangements at the *Bcl11b* locus. The previously described intron 1 and intron 3 cRSSs are indicated (*; Sakata *et al.*; Champagne *et al.*). (B) Estimated recombination frequency of the previously described *Bcl11b* intron 1 and intron 3 cRSS pair. The gel shows 25 nested-PCR reactions containing 100 ng of WT total-thymocyte gDNA per reaction, corresponding to approximately 417,000 cells in total. Eleven recombination events were detected, giving an estimated frequency of 26.4 rearrangements per million cells, consistent with previously reported frequencies. (C) Agarose gel images showing rearrangements involving the newly identified cryptic sites III and IV and the *Bcl11b* intron 3 cRSS partner. PCR products were sequenced to confirm the identity of the detected rearrangements.

**Figure S4.**
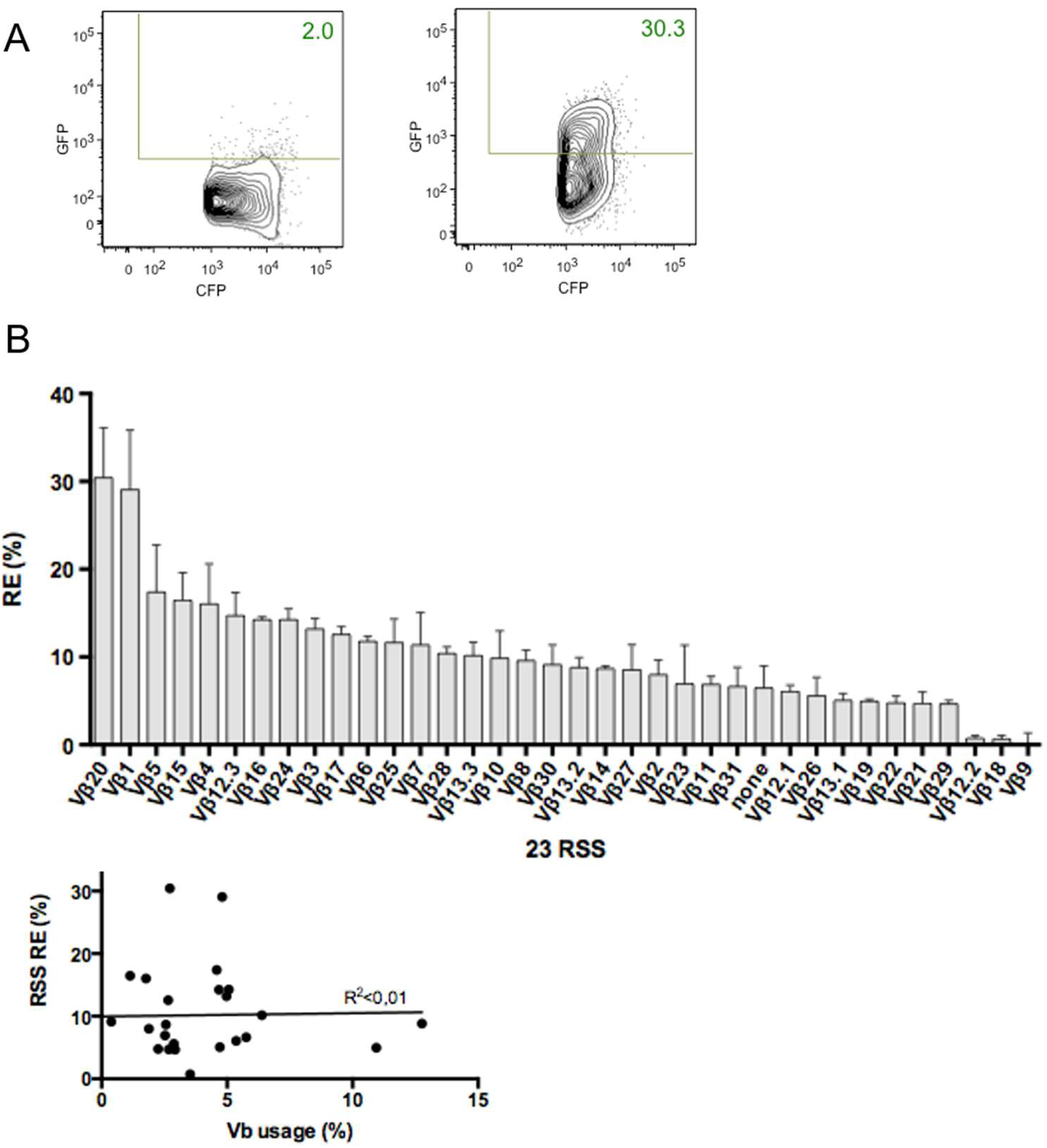
***In vitro* RSS recombination efficiency is poorly associated with gene-segment usage *in vivo*.** (A) Representative flow-cytometry plots of Vβ1–Dβ2 recombination in 293T cells co-transfected with the GFPi reporter construct in the absence (left) or presence (right) of RAG-expression plasmids. (B) *In vitro* recombination efficiency of all Vβ RSSs paired with either the Dβ2 12-RSS or the consensus 12-RSS (ConS12) (see Table S6 and Trancoso *et al.*). A construct containing the 12-RSS without a 23-RSS partner (“none”) was included as a negative control. Recombination efficiency (RE) is shown for each RSS pair as the mean ± SD from at least three independent experiments. RSS pairs are ordered according to their RE when paired with Dβ2. (C) Relationship between the *in vitro* recombination efficiency of each TCRβ Vβ RSS and the *in vivo* usage of the corresponding gene segment in DN3 thymocytes, as reported by Gopalakrishnan *et al.* No clear association was observed, as indicated by the low coefficients of determination (R²).

**Figure S5.**
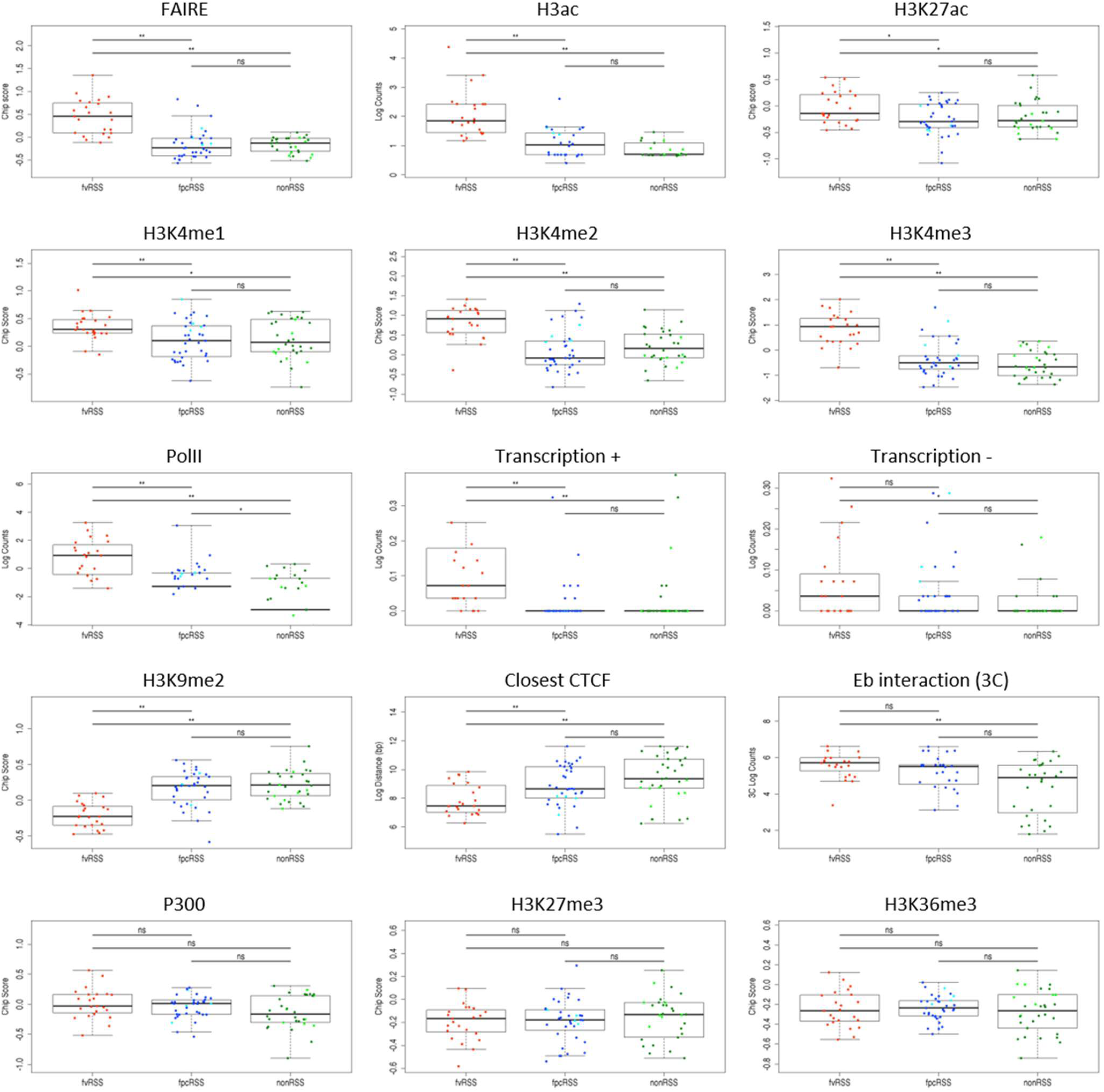
Comparison of genetic and epigenetic features among recombination-site groups. Boxplots show features associated with chromatin accessibility, histone modifications, transcription and chromatin organization for functional Vβ RSSs (vRSSs), sites with detected cryptic recombination (cRSSs) and tested sites at which recombination was not detected (nonRSSs). Statistical comparisons between groups are indicated above each plot. Several features distinguish functional Vβ RSSs from one or both of the other groups, whereas none reliably distinguishes cRSSs from nonRSSs.

**Figure S6.**
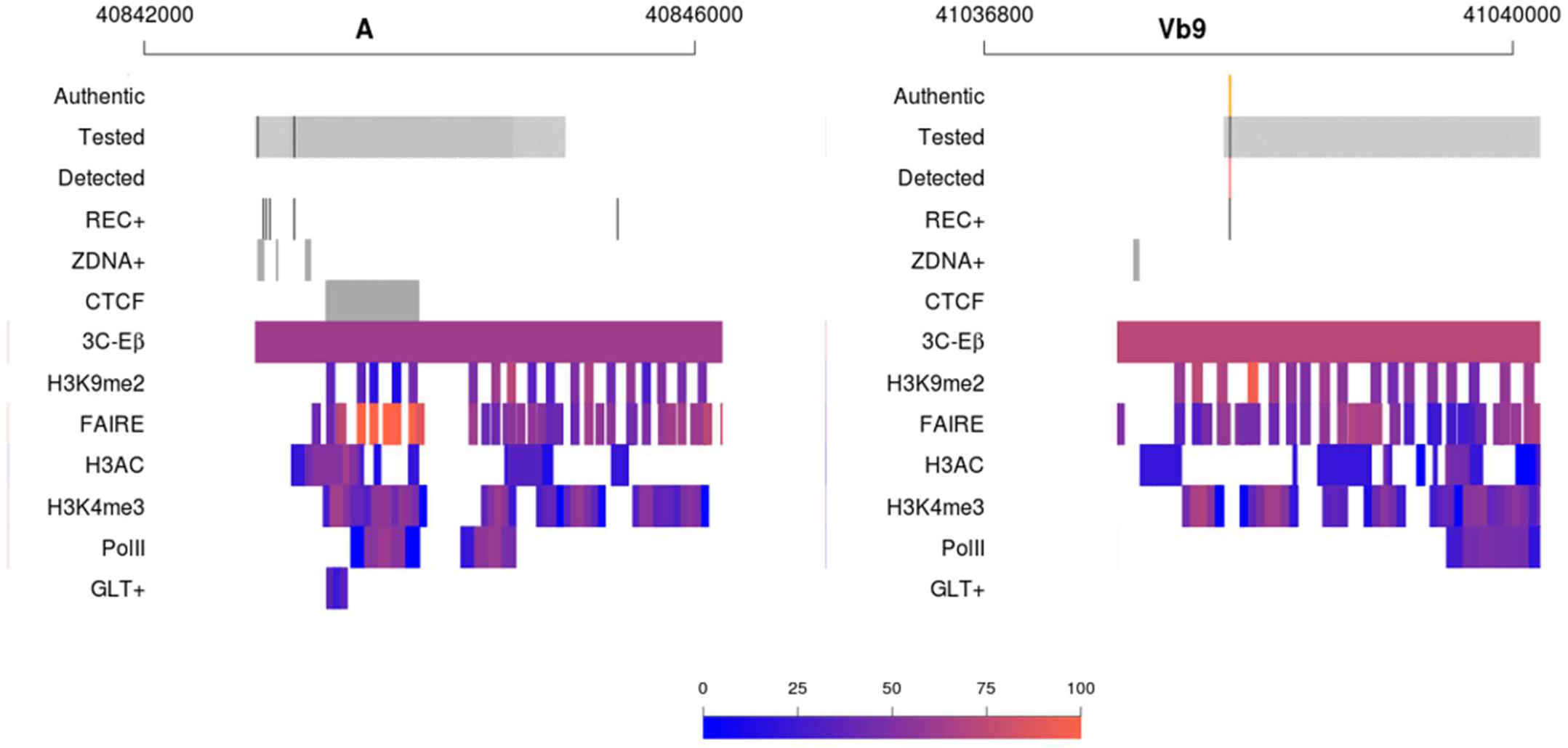
Examples of genetic and epigenetic profiles at sites with and without detectable recombination. A and Vβ11 (top) are examples of tested sites at which no rearrangement was detected despite local genetic and epigenetic features associated with recombination-permissive conditions. Conversely, rearrangements were detected at Vβ9 and Z43 (bottom) despite comparatively low levels of epigenetic marks associated with accessible chromatin.

**Figure S7.**
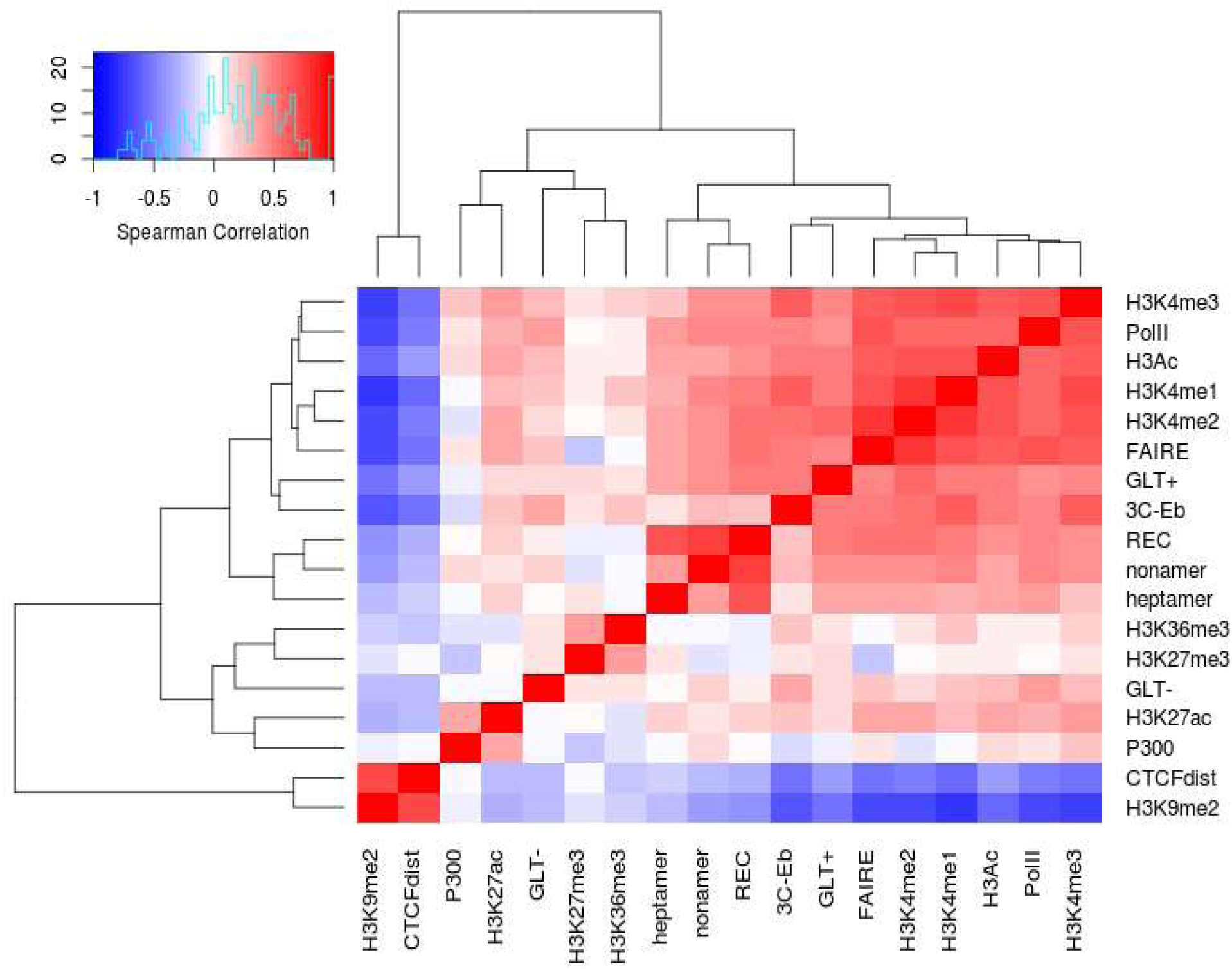
Correlations among genetic, epigenetic and chromatin-organization features at the TCRβ locus. The matrix shows pairwise correlations among the features analysed, including histone modifications (H3K4me1, H3K4me2, H3K4me3, H3ac, H3K27ac, H3K27me3 and H3K36me3), transcription-associated features (positive- and negative-strand germline transcription [GLT+ and GLT−], Pol II and P300), chromatin accessibility (FAIRE), chromatin organization (CTCF and 3C-Eβ), and the REC score.

### SUPPLEMENTARY TABLES

**Table S1.**
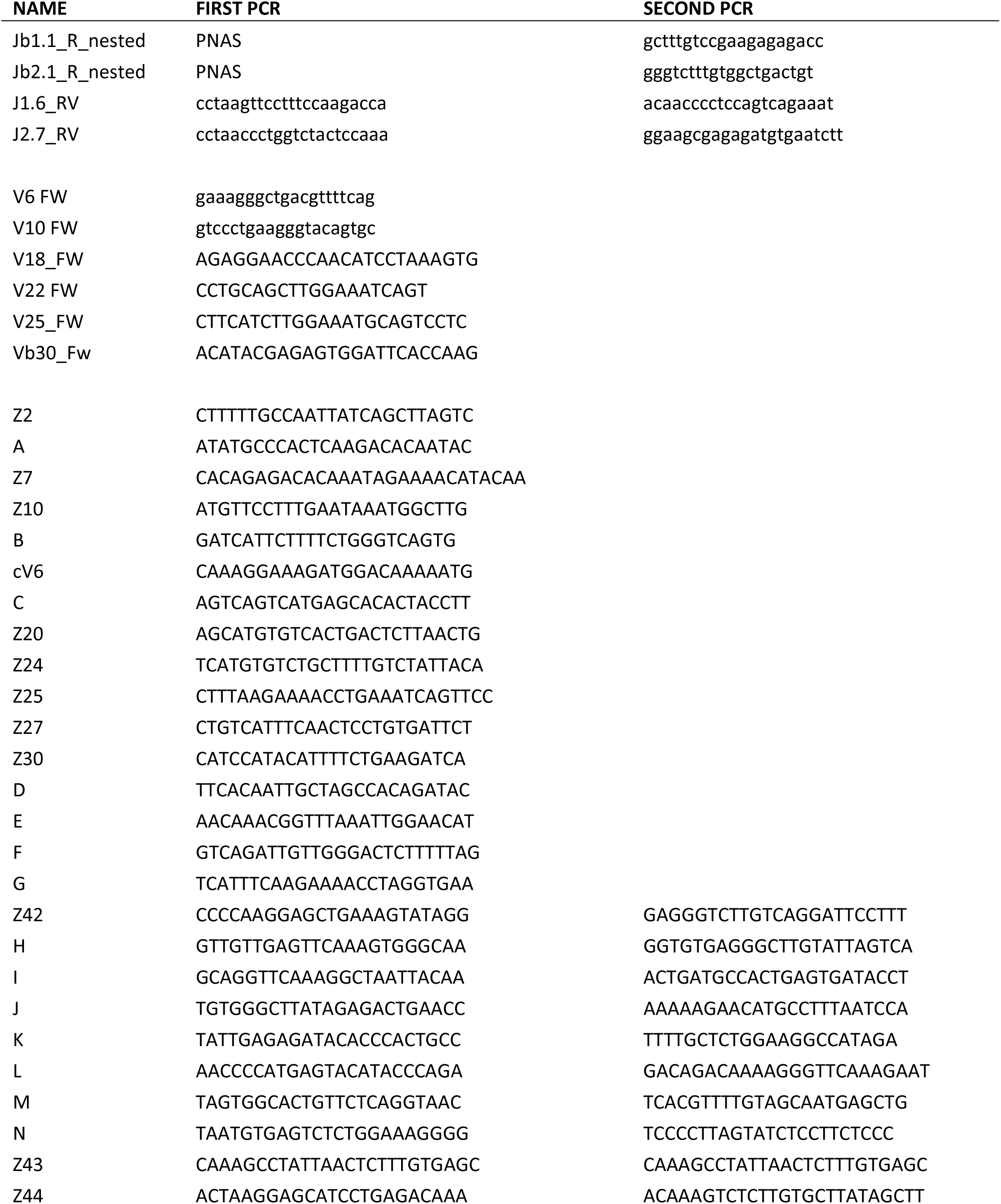
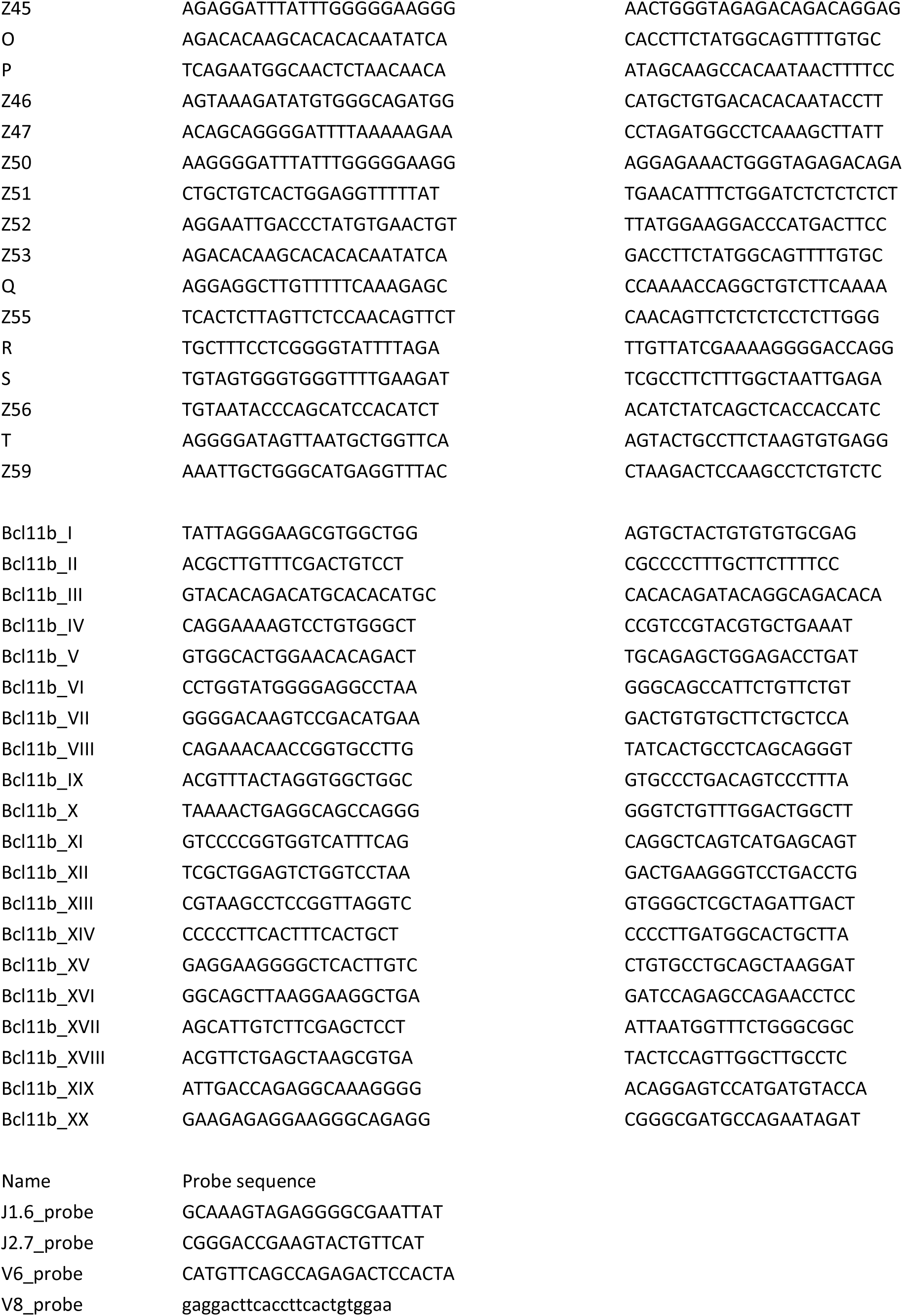
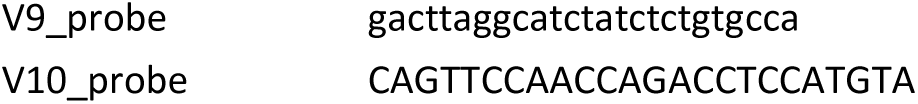
Primers and probes for (semi-)nested PCR assays.

**Table S2.**
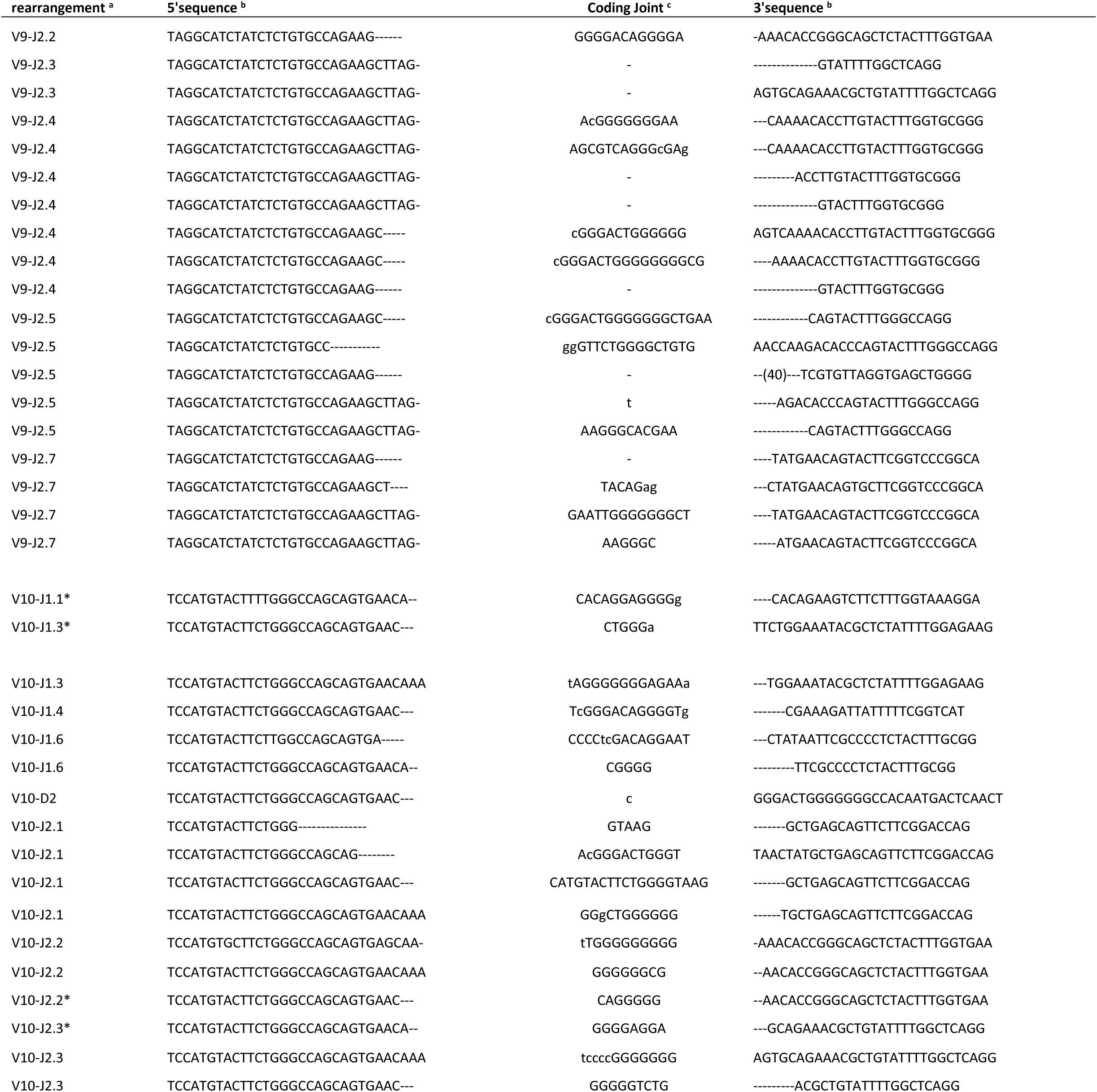
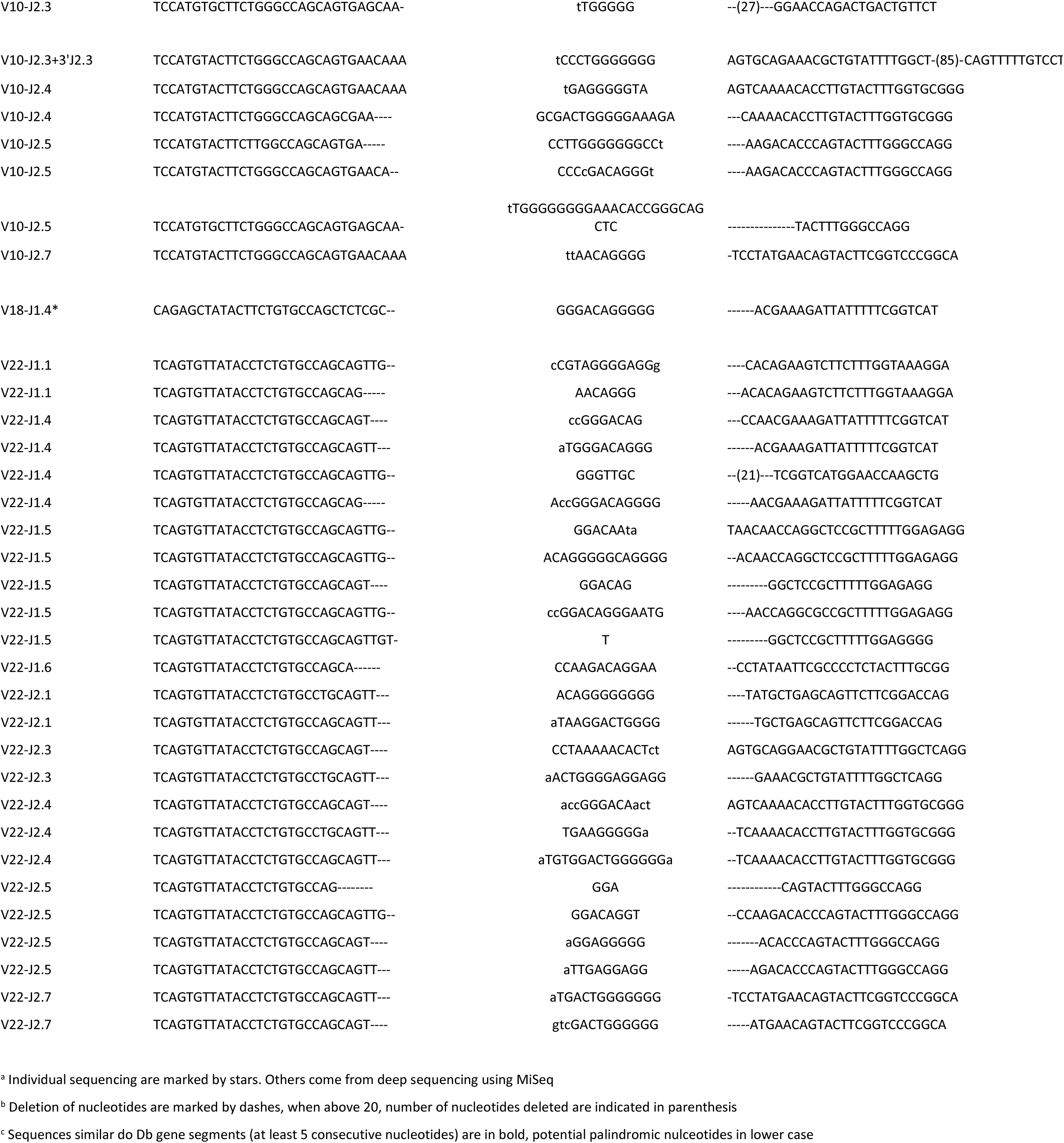
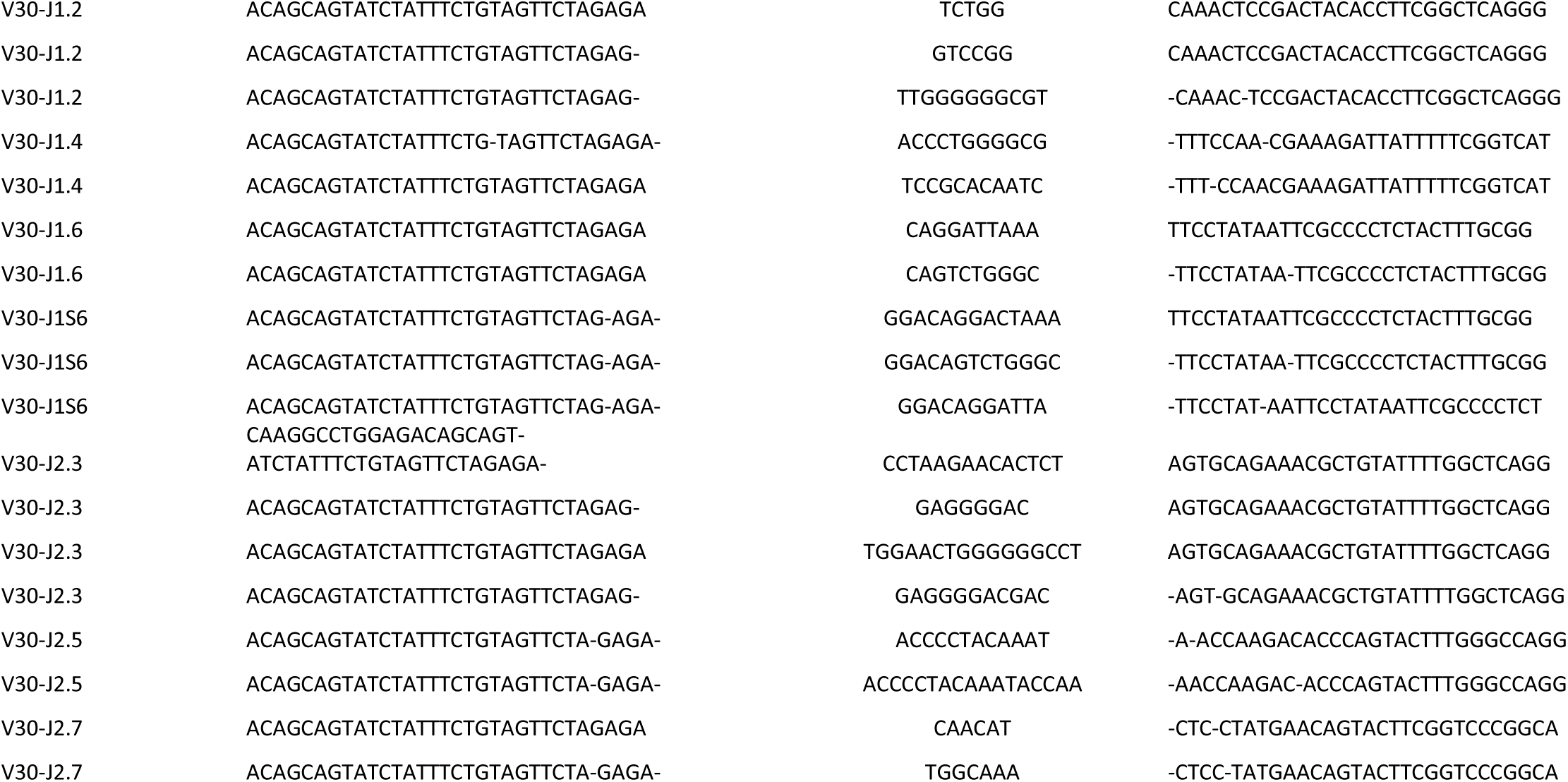
Pseudogene rearrangements coding joints.

**Table S3.**
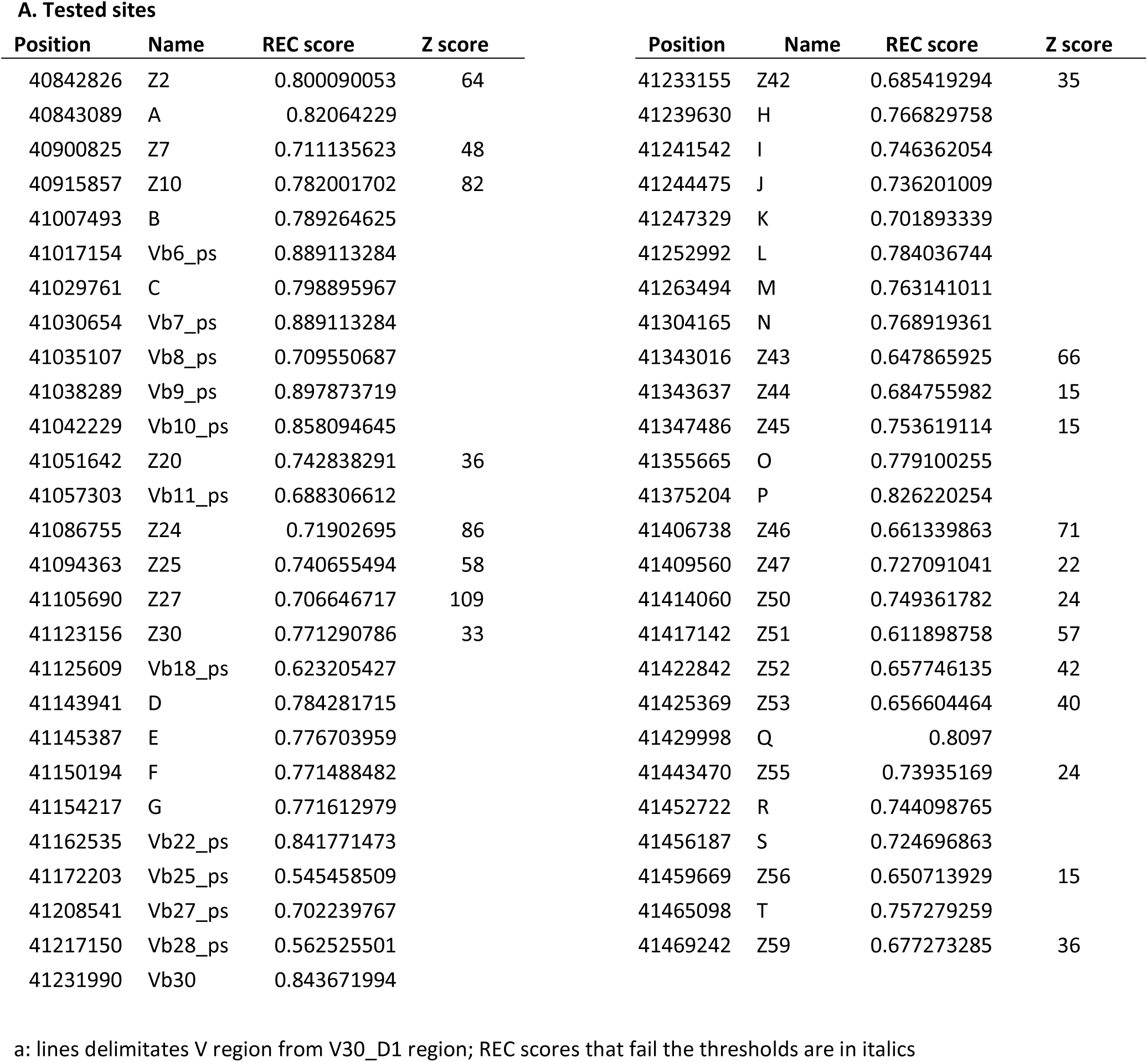
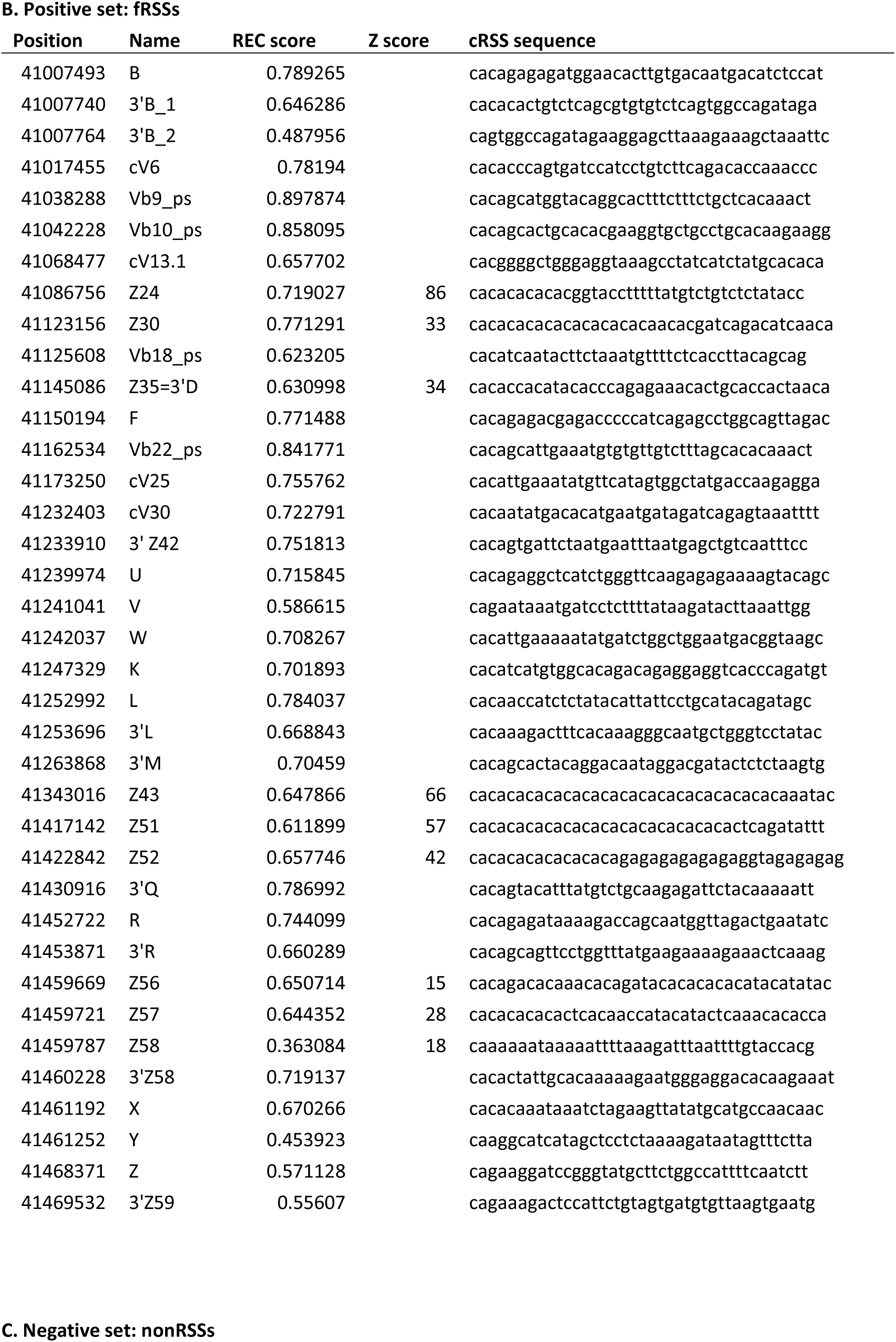
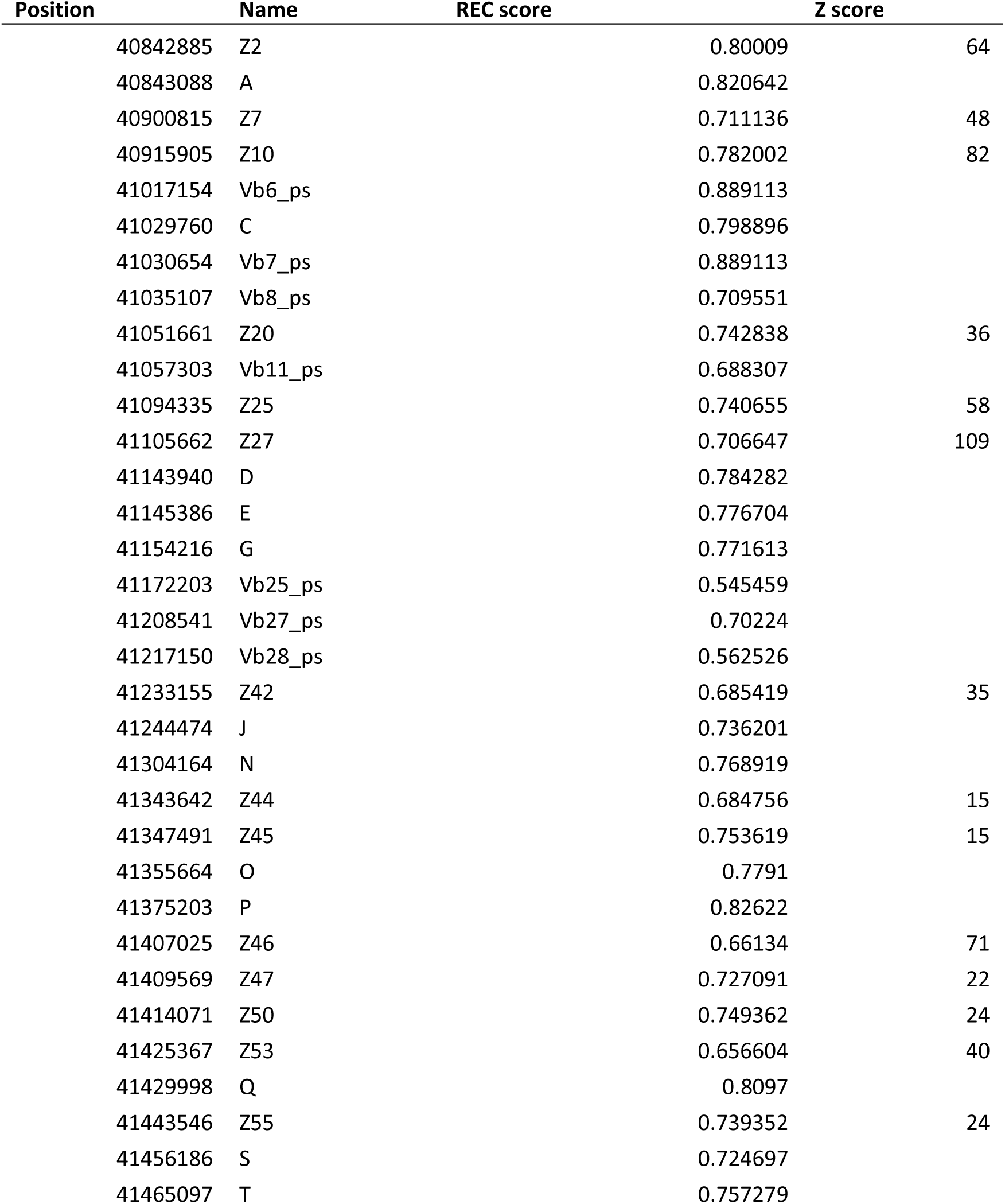
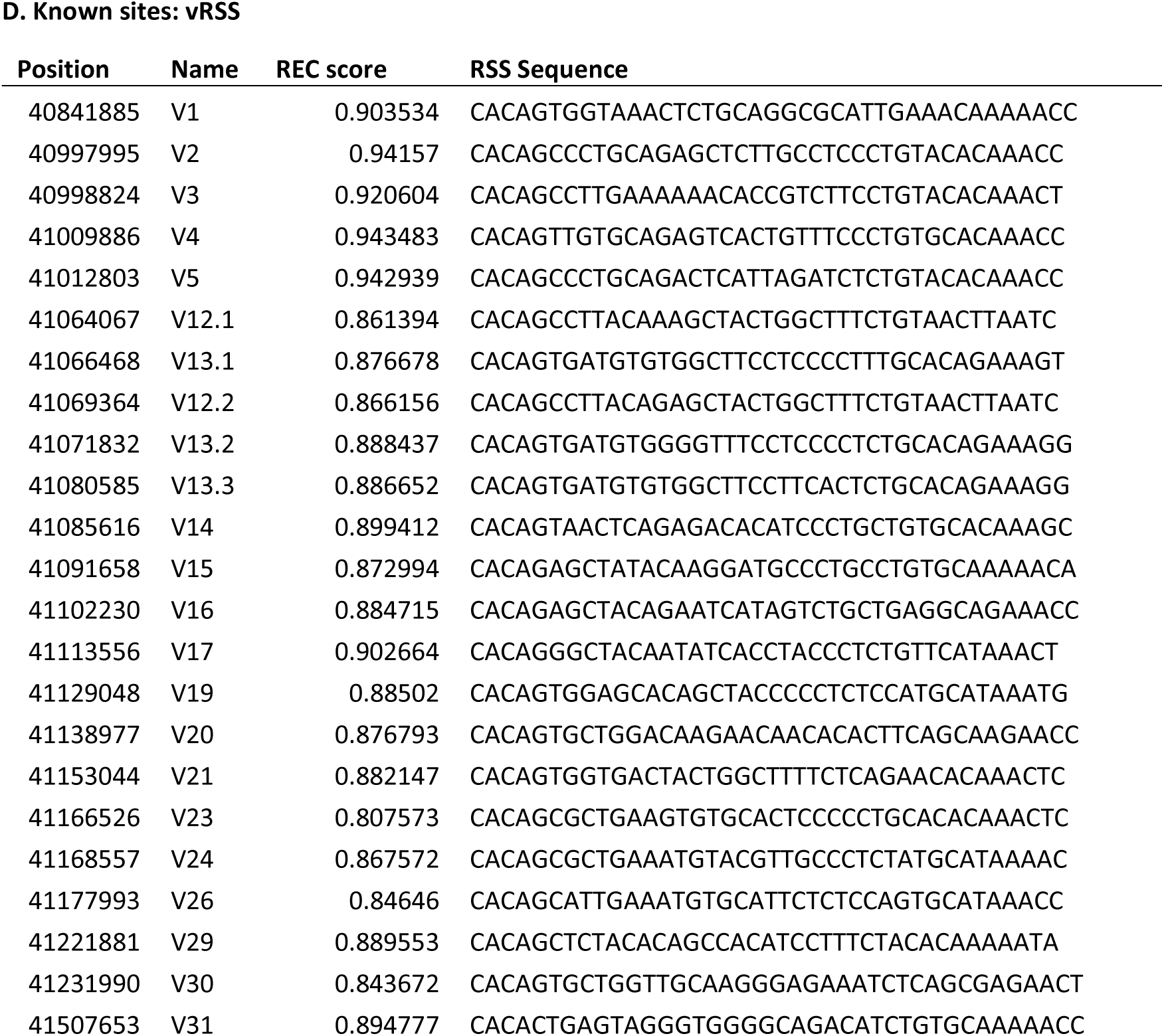
Coordinates, REC and Z-scores of tested sites by PCR (A), positive sites (B), negative sites (C) from deep-sequencing results a, (D) known functional sites.

**Table S4.**
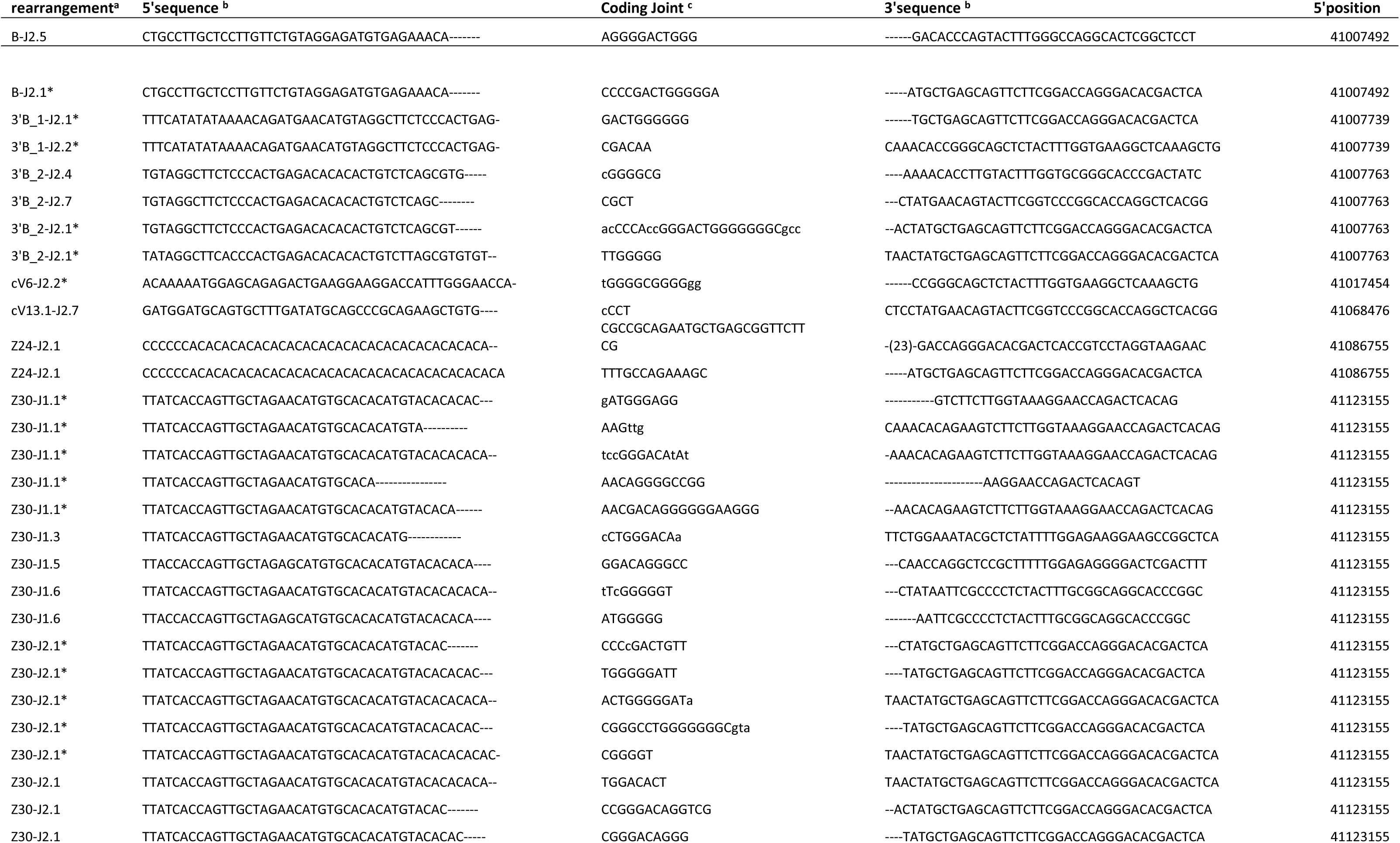
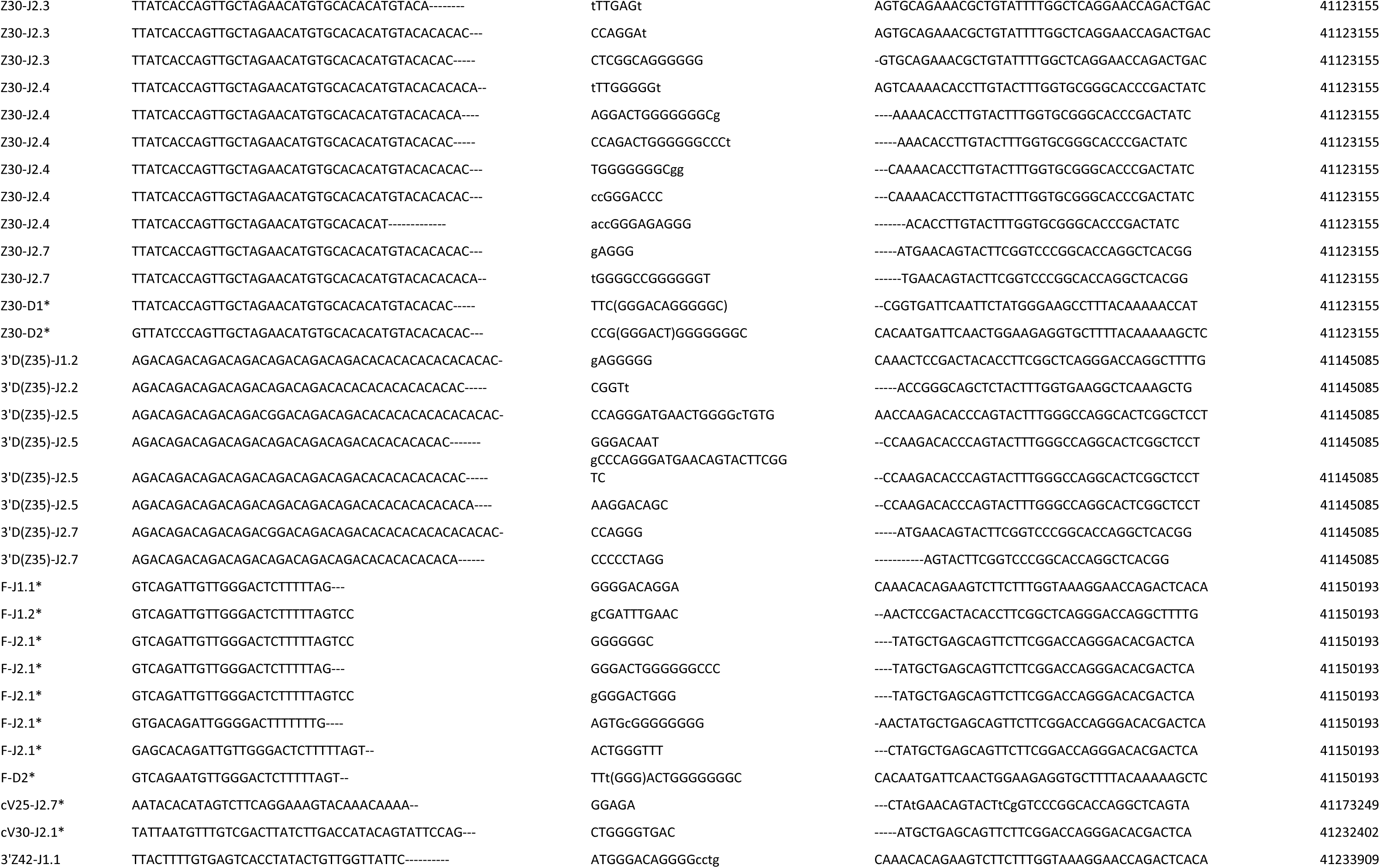
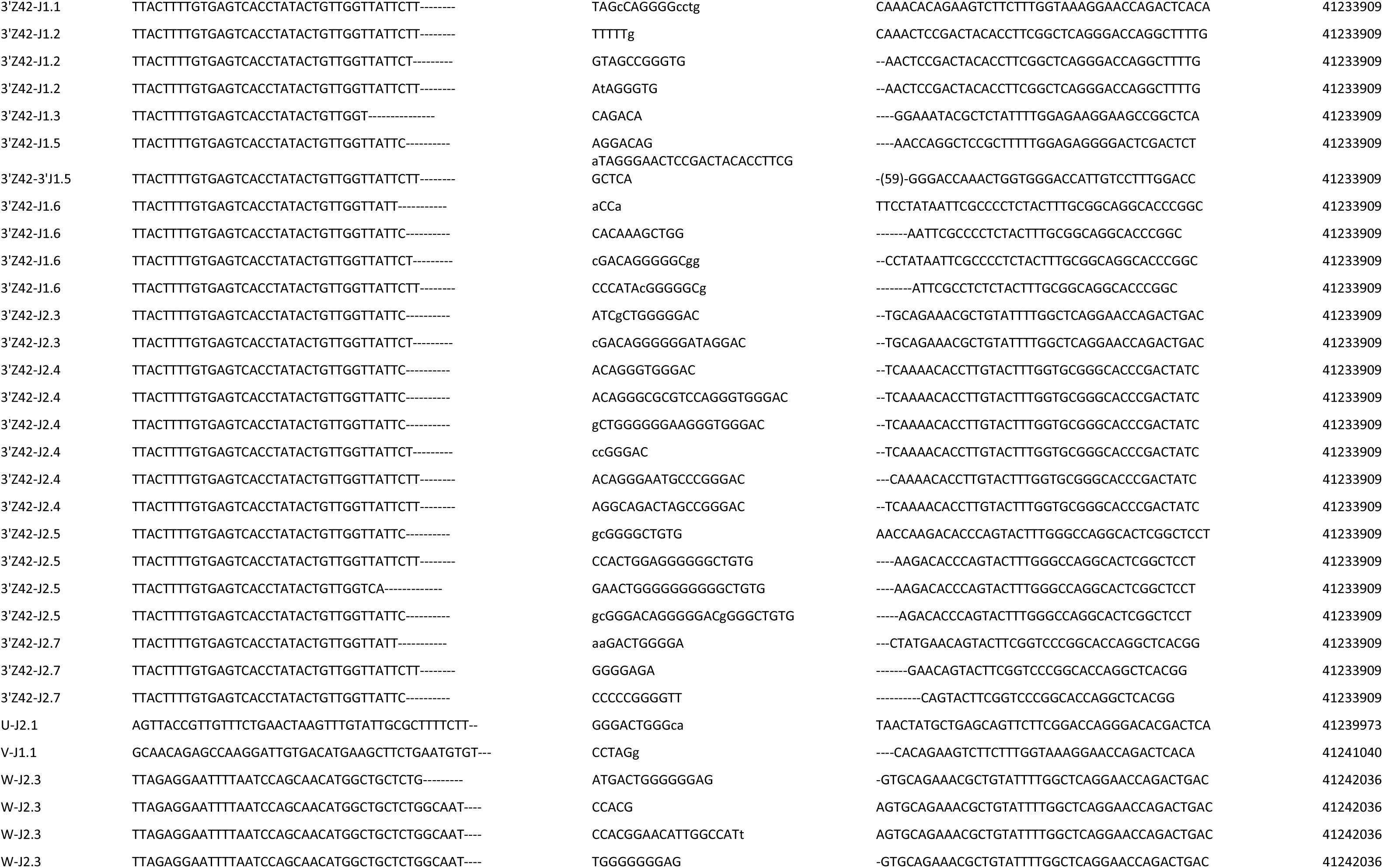
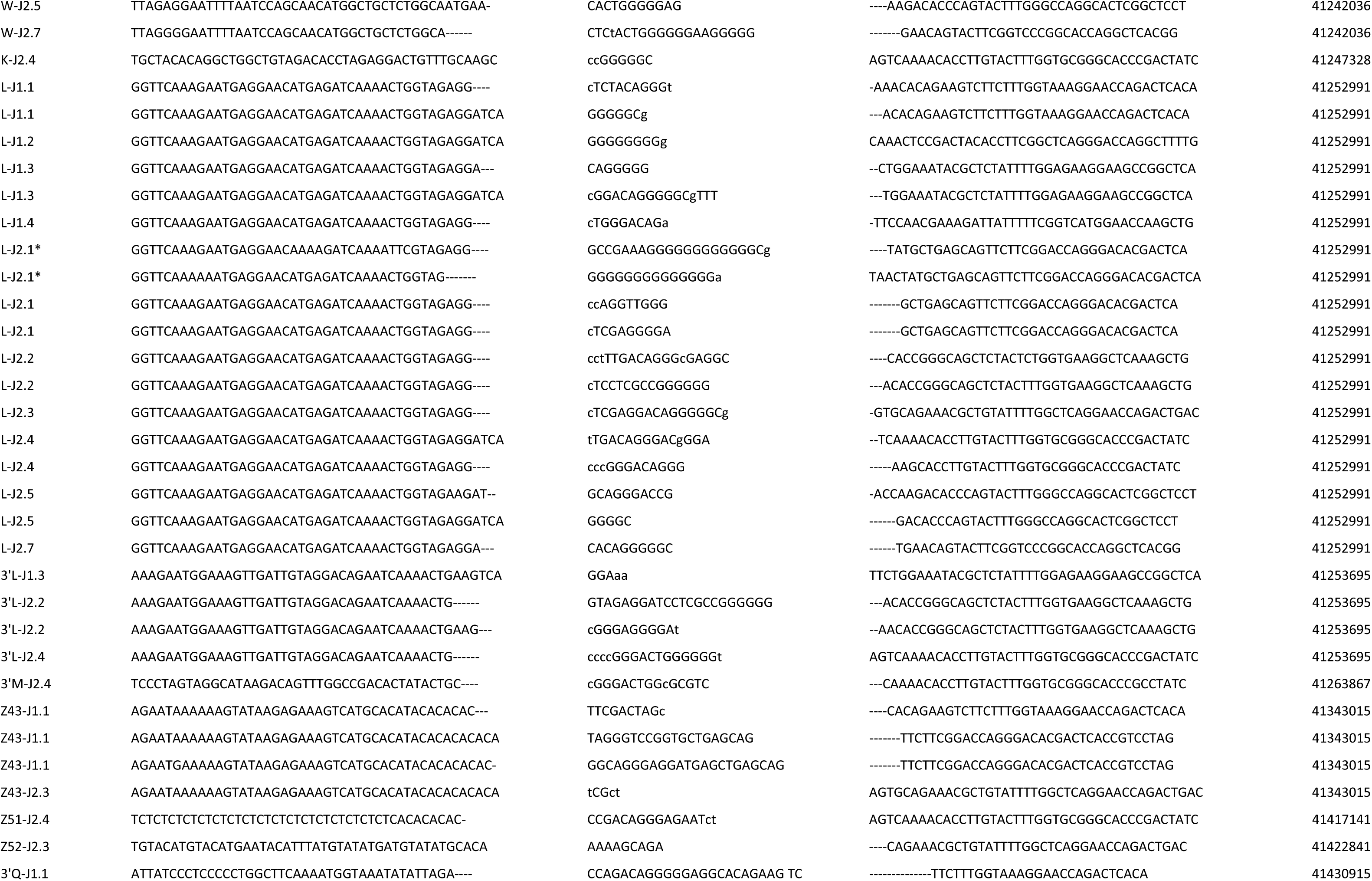
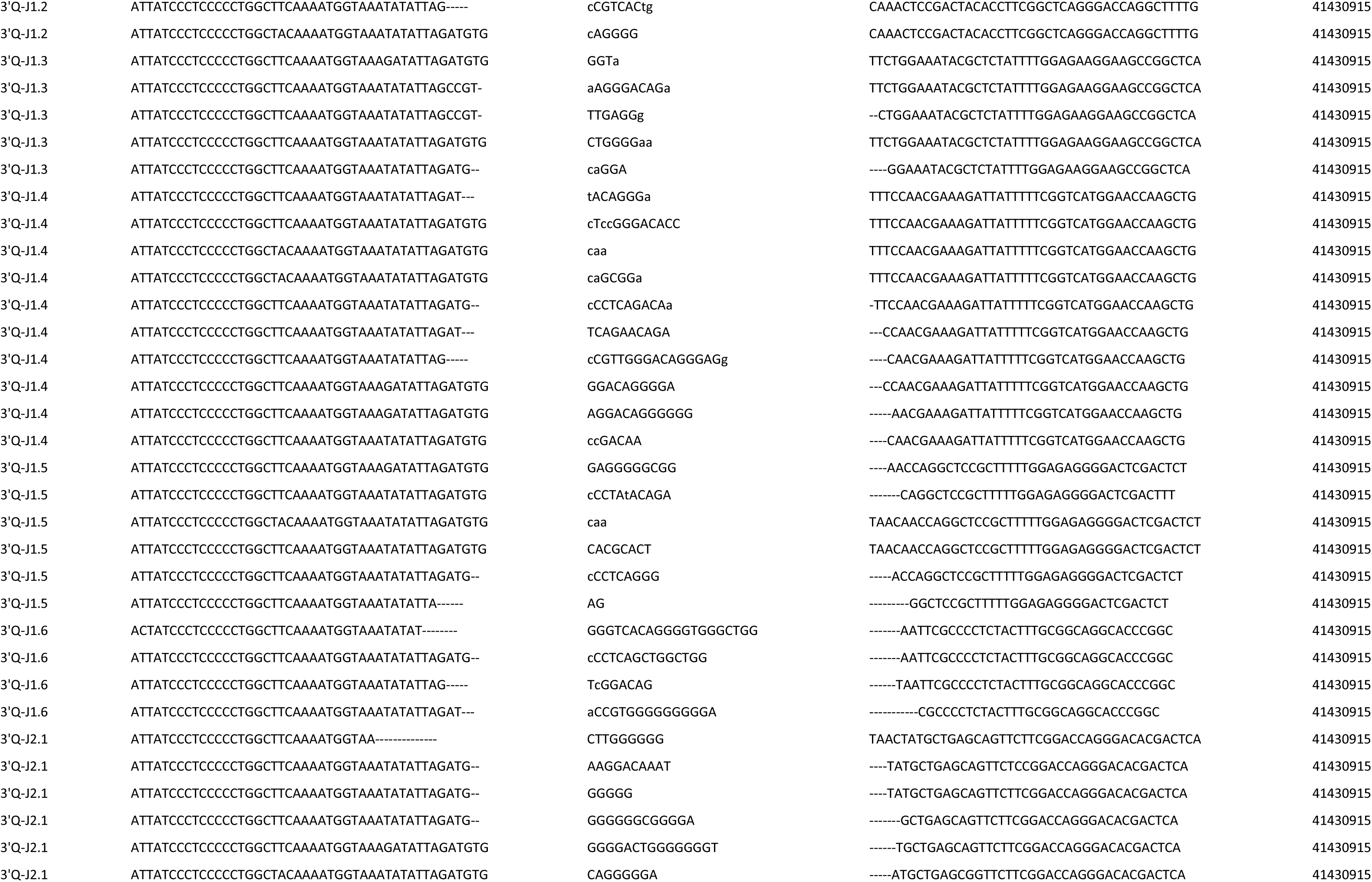
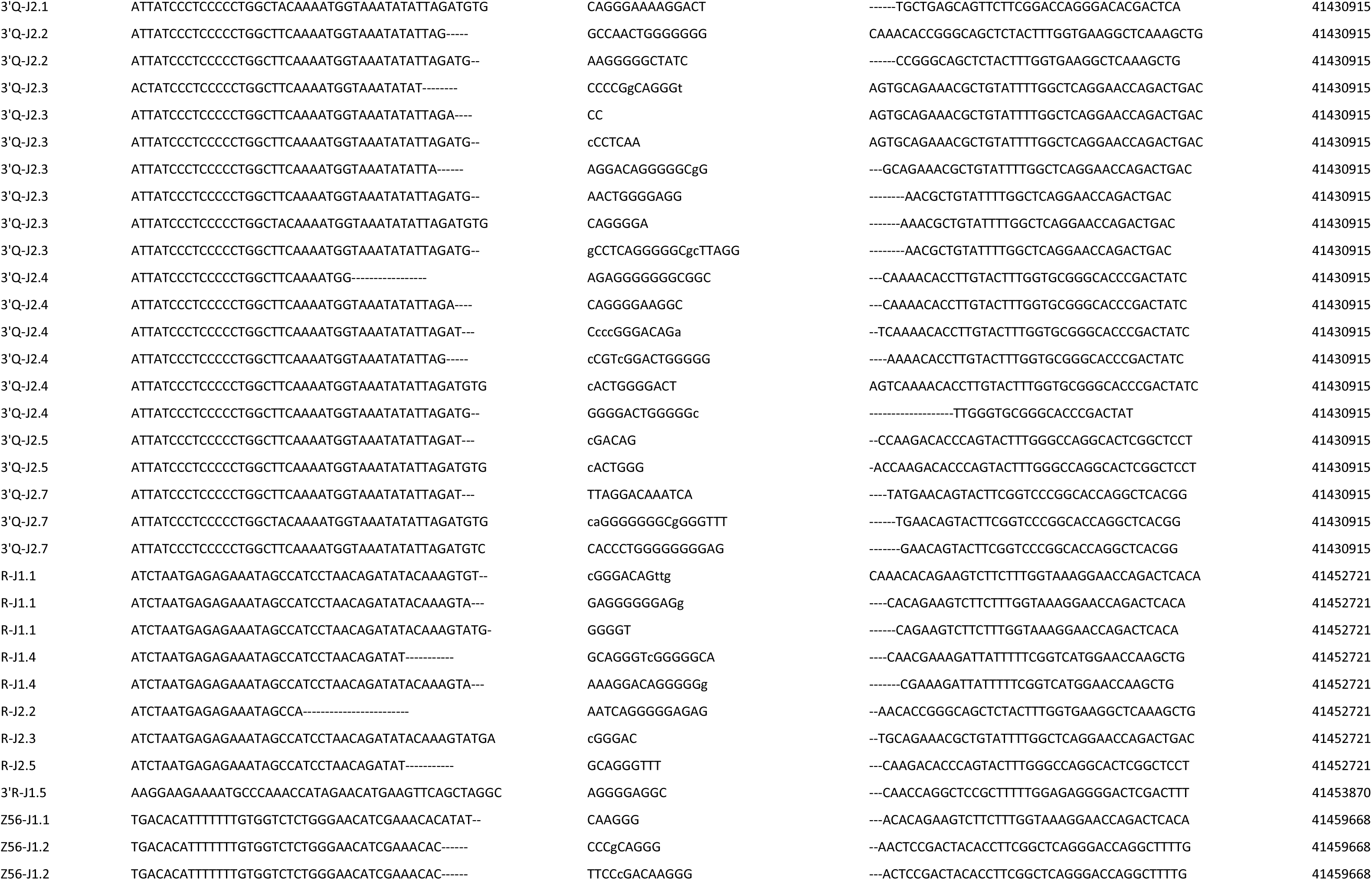
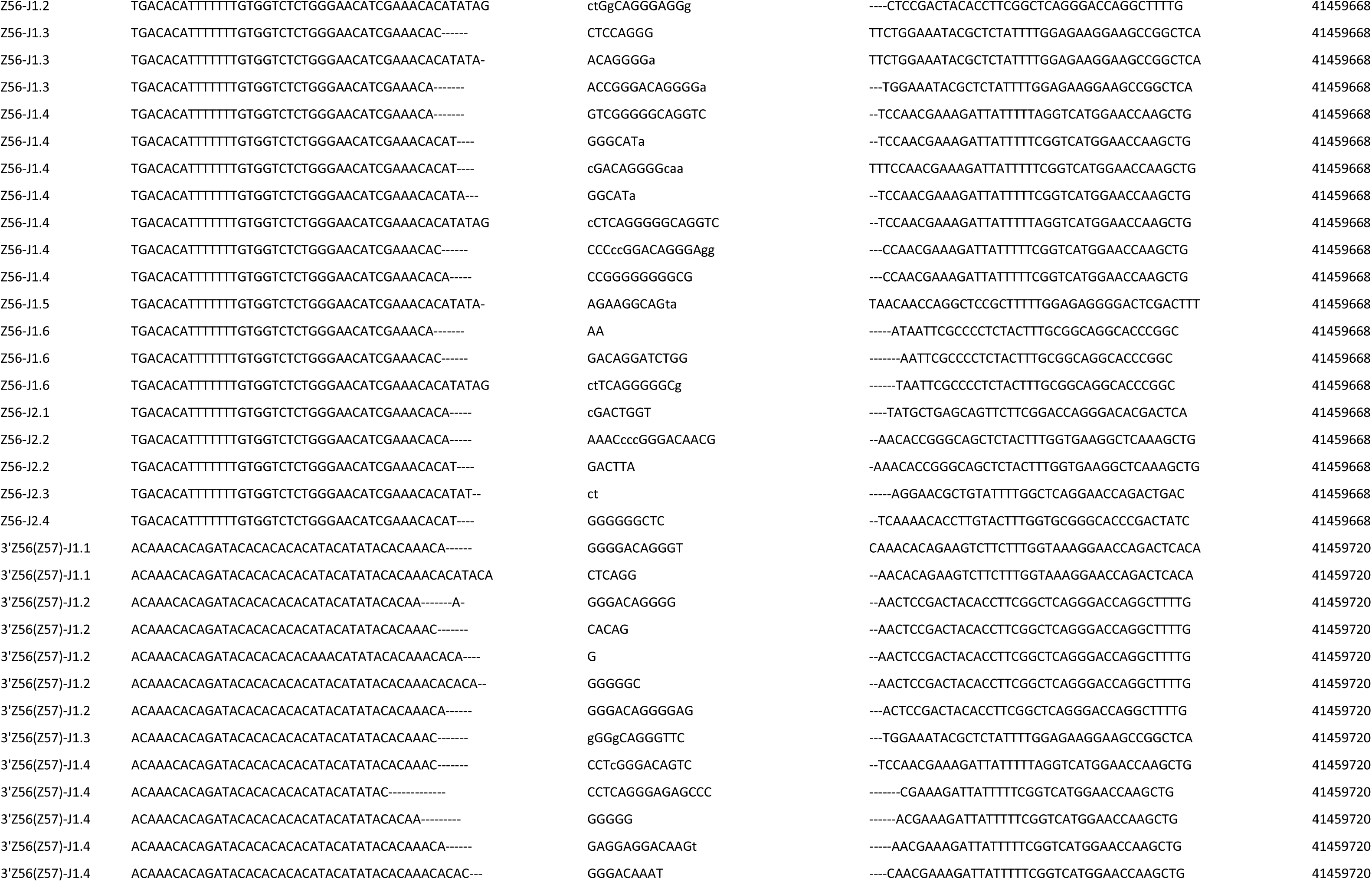
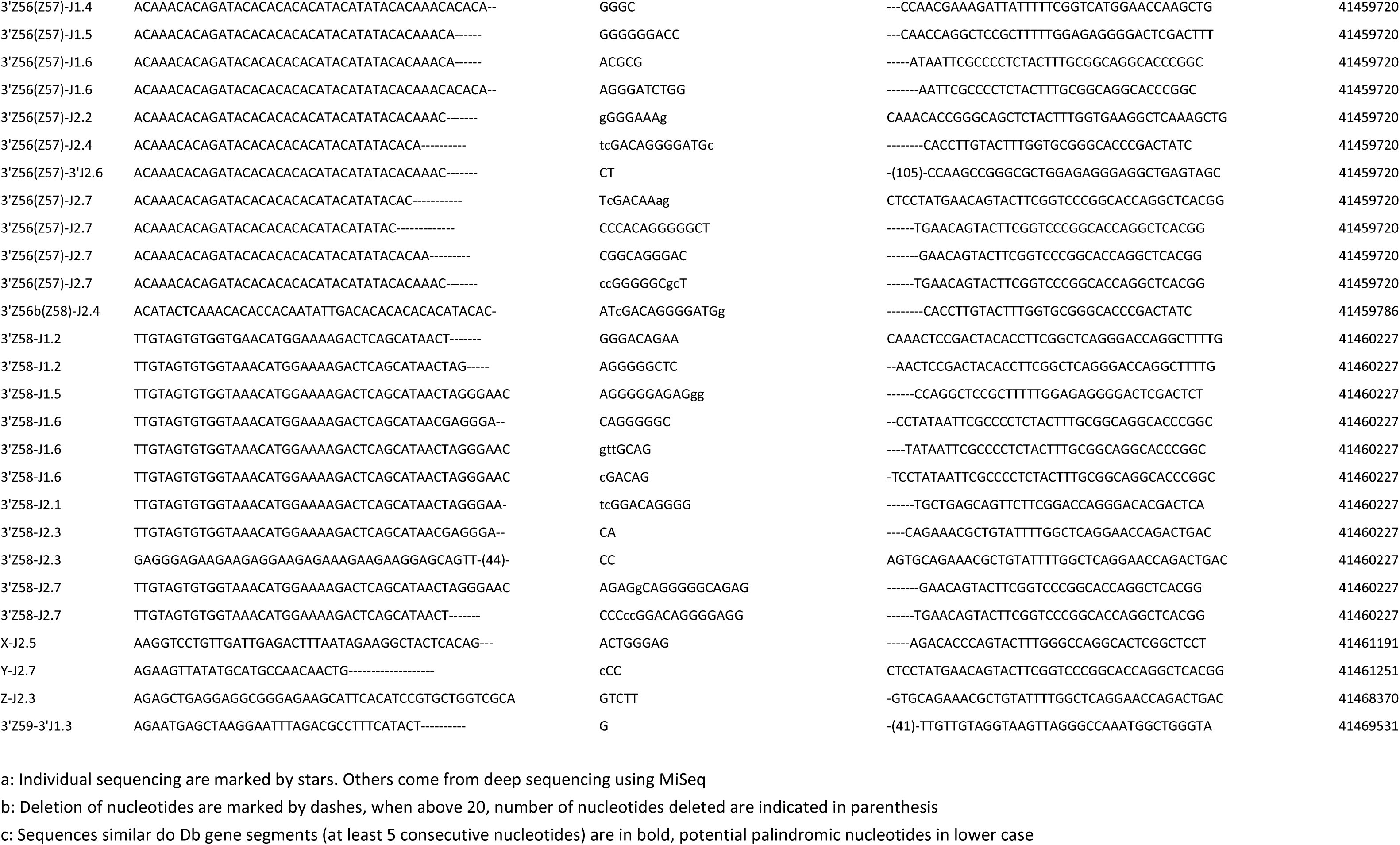
Cryptic sites rearrangements coding joints.

**Table S5.** Sites tested by PCR at the Bcl11b gene.

| **Name** | **Site position** | **REC score\*** | **Z score\*** |
| --- | --- | --- | --- |
| Bcl11b_I | 109169382 | 0.37802006 | 57 |
| Bcl11b_II | 109171408 | 0.73304724 |  |
| Bcl11b_III | 109176788 | 0.7786535 | 27 |
| Bcl11b_IV | 109177252 | 0.72159439 | 301 |
| Bcl11b_V | 109183231 | 0.76938862 | 15 |
| Bcl11b_VI | 109183943 | 0.74628513 |  |
| Bcl11b_VII | 109187690 | 0.59032114 | 29 |
| Bcl11b_VIII | 109205324 | 0.74740807 |  |
| Bcl11b_IX | 109207034 | 0.73620101 | 46 |
| Bcl11b_X | 109207567 | 0.47601419 | 24 |
| Bcl11b_XI | 109210517 | 0.73691734 |  |
| Bcl11b_XII | 109211960 | 0.42645711 | 77 |
| Bcl11b_XIII | 109213335 | 0.63843253 | 26 |
| Bcl11b_XIV | 109216167 | 0.76065402 |  |
| Bcl11b_XV | 109224131 | 0.45036689 | 37 |
| Bcl11b_XVI | 109228276 | 0.45037033 | 55 |
| Bcl11b_XVII | 109229701 | 0.59840657 | 35 |
| Bcl11b_XVIII | 109230922 | 0.74691381 |  |
| Bcl11b_XIX | 109232576 | 0.45895961 | 46 |
| Bcl11b_XX | 109241282 | 0.45862291 | 90 |

\* best REC and Z scores within 300bp downstream of the primer used

**Table S6.**
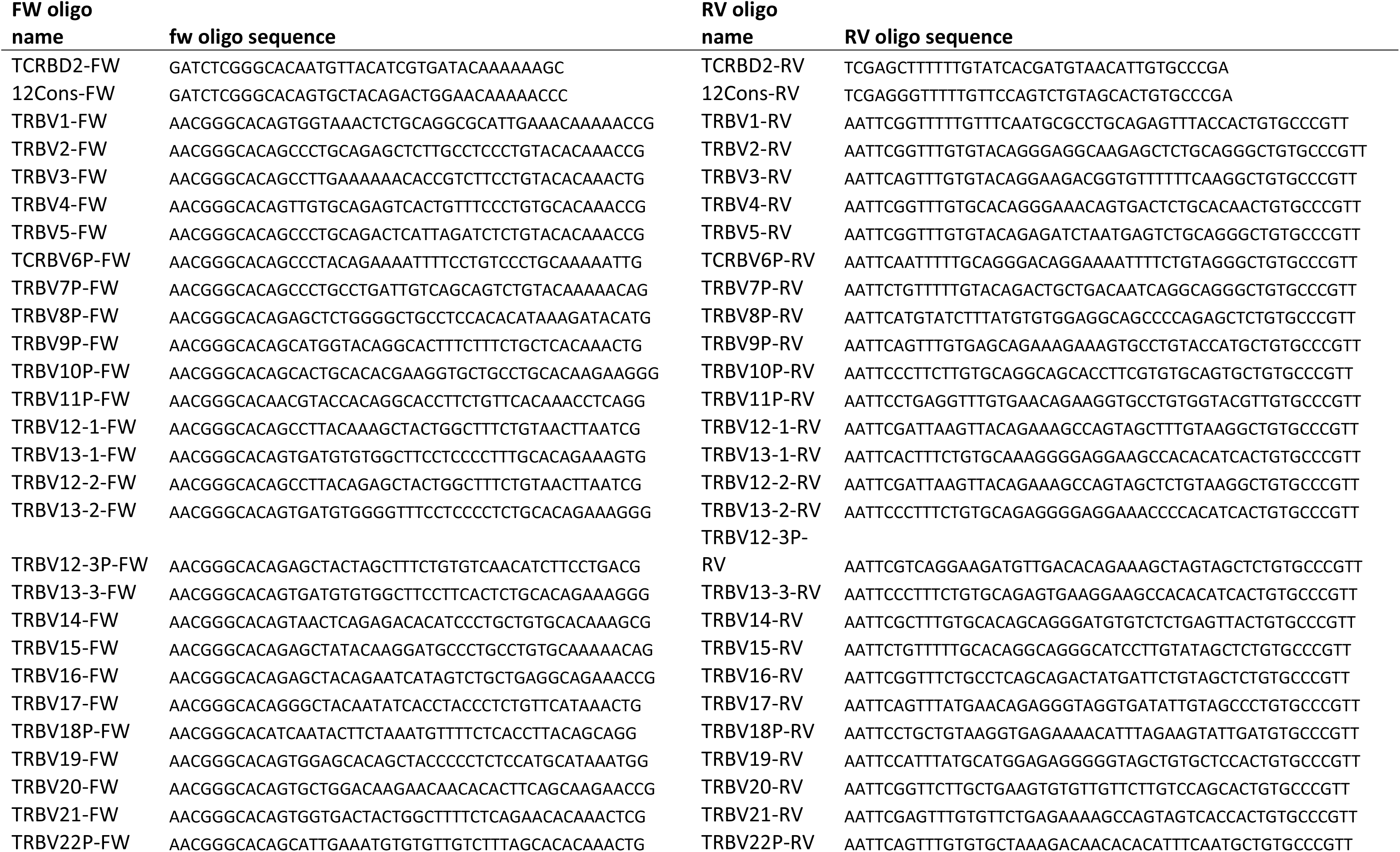
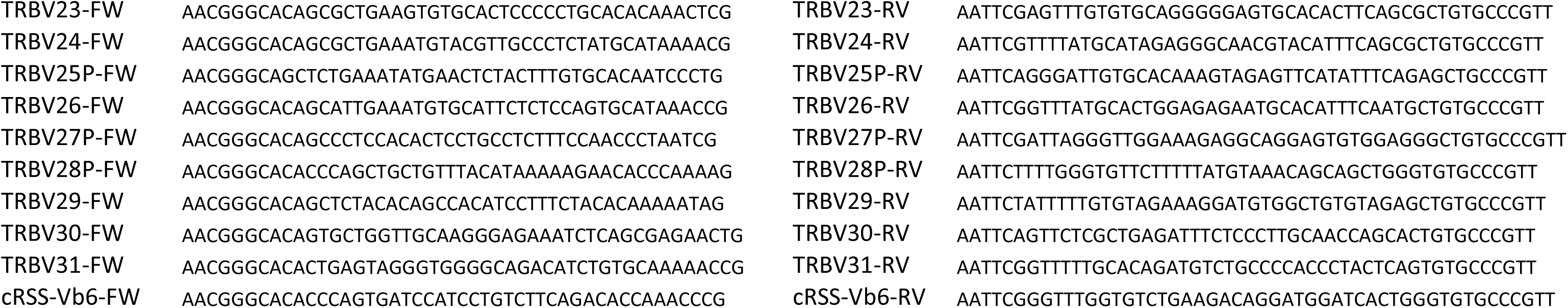
Oligonucleotides for RSS cloning in the GFPi assay.

